# Reducing work-load of pathotype and gene detection in *Klebsiella pneumoniae* by leveraging machine learning

**DOI:** 10.1101/2023.10.02.560438

**Authors:** Rahimeh Sanikhani, Kourosh Alizadeh Kiani, Mehdi Soroush, Mohammad Moeinirad, Seyed Ahmad Sanikhani, Masoumeh Beheshti, Sajad Tavakoli, Farzad Badmasti, Seyed Hossein Sadati

**Affiliations:** Department of Bacteriology, Pasteur Institute of Iran, Tehran, Iran; Department of Mechanical Engineering, K. N. Toosi University of Technology, Tehran 19919-43344, Iran; Division of Bacteriology, Department of Pathobiology, School of Public Health, Tehran University of Medical Sciences, Tehran, Iran; Department of Mechanical Engineering, Amirkabir University of Technology (Tehran Polytechnic), Tehran 15875-4413, Iran; Pathophysiology laboratory, Sina hospital, Tehran University of Medical Sciences, Tehran, Iran; Department of Biotechnology and Biomedicine, Technical University of Denmark, Kongens Lyngby, Denmark; Microbiology research center (MRC), Pasteur Institute of Iran, Tehran, Iran

**Keywords:** *Klebsiella pneumoniae*, beta-lactamase genes, PCR, pathotypes, Machine learning

## Abstract

**Background:** The growing prevalence of carbapenem resistance has caused an increasing number of bacterial isolates with multi-drug resistance features, especially in the *Enterobacteriaceae* family. *Klebsiella pneumoniae*, as one of the important members of the *Enterobacteriaceae* family, causes serious infections, which has attracted the attention of scientists due to the emergence of hypervirulent pathotypes with increasing antibiotic resistance and has been raised as a major concern worldwide. Early detection of this new super bacterium and its antibiotic resistance is of great help in reducing mortality and costs. The lack of new antibiotic options underscores the need to optimize current diagnostics. Therefore, this study was designed to leverage machine-learning approach for optimized selection of crucial antibiotics to reduce the experiments needed for the detection of pathotypes and genes’ presence in two classical and hypervirulent *K. pneumoniae* pathotypes.

**Methods:** 341 non-duplicate clinical isolates of *K. pneumoniae* were collected from five university hospitals in Tehran and Qazvin, Iran. Pathotype differentiation of classical (c*Kp*) and hypervirulent *K*.*pneumoniae* (hv*Kp*) was done by PCR method by two molecular biomarkers including *iuc* and *iut*. After identifying the phenotypic antibiotic resistance, the presence of antibiotic resistance genes was detected by PCR method. Then, the relevance of resistance/susceptibility of the antibiotics and presence of pathotypes, aerobactin, and beta-lactamase genes was investigated and analyzed using five supervised machine learning algorithms by selecting crucial antibiotics through feature selection methods.

**Results:** Among the 341 *K*.*pneumoniae* isolates, 102 and 239 isolates were hv*Kp* and c*Kp* respectively. The highest rate of antibiotic resistance after ampicillin (100%) was related to cefotaxime (76.2%) and the lowest rate of resistance was found in meropenem (24.3%). Imipenem, Meropenem, Aztreonam, Ceftazidime, Ceftriaxone, and Gentamicin are crucial antibiotics for detection of the pathotypes and the aerobactin genes. Moreover, Cefotaxime, Ciprofloxacin, Cefepime, Meropenem, and Imipenem are essential for detection of the beta-lactamase genes.

**Conclusion:** Implementing a machine learning approach including various feature selection methods and algorithms, results in less-required experiments on more limited antibiotics to detect genes and pathotypes. Our findings reveal that using machine learning in the prediction of the presence of genes and pathotypes of clinical isolates was a suitable method in terms of rapidity and cost-efficiency on top of accuracy.

## 1. Introduction

*Klebsiella pneumoniae* is a member of the *Enterobacteriaceae* family and one of the most common opportunistic pathogens among enteric gram-negative bacilli (1, 2). It is known as commensal bacteria in animal/human normal intestinal microflora. Therefore, they can be found in the environment, soil, sewage, and polluted food sources, causing various diseases including urinary tract infections, respiratory infections, sepsis, and wound infections (3, 4). A common phenomenon in *Enterobacteriaceae* is resistance to different antibiotic classes, called multi-drug resistance (MDR). In recent years, the increase in infections caused by MDR *K. pneumoniae* due to the inappropriate use of antibiotics in the bacterial infections treatment protocol has caused public health threat and burdened high mortality(5). Recent studies based on genetic characteristics have defined three groups of *K*.*pneumoniae* strains, which include: opportunistic or classical (c*Kp*), hypervirulent (hv*Kp*), and multi-drug resistant (MDR-*Kp*) *K*.*pneumoniae* (6). The hv*Kp*s are a potential pathogen due to the large virulence plasmid and have lower antibiotic resistance than c*Kp*. However, in recent years, we have observed an increase in the community-acquired infections caused by MDR-hv*Kp*s (7, 8). The high-risk pathogenic clones of hvK*p* that are antibiotic resistant could be counted as a significant global threat (9, 10). Therefore, rapid detection of these strains, so-called “superbug,” is crucial.

Diagnostic tests are a core component of modern healthcare practice. High-quality diagnostics have become increasingly essential, especially in the light of rising multi-drug resistance. However, providing information as the basis for infectious disease management is complex. Antimicrobial susceptibility testing (AST) has experienced little change over the years. It still relies on culture-dependent methods; consequently, clinical microbiology diagnostics is labor-intensive, expensive, and slow. Therefore, these limitations underscore the need for an optimized method in order to select the most influential antimicrobial drugs.

In the last decade, machine learning-based (ML) approaches have received increasing attention especially in biology, and many researches based on ML have recently been conducted (11-13). This study employs a machine learning approach in a binary classification problem to circumvent the mentioned gaps (reducing work-load in the experiments needed for the detection of pathotypes and genes’ presence). Detections of the presence of the genes and the pathotypes were investigated through selection of crucial antibiotics and implementing them in ML algorithms as input variables. For this purpose, different feature selection methods were employed to designate crucial antibiotics. Then, five supervised ML algorithms were separately trained based on the selected features (isolates’ resistance/susceptibility to the antibiotics). We built predictive models to identify crucial antibiotics for detection of pathotypes and genes’ presence. Our proposed method leads to the selection of the most efficient antibiotics that significantly reduces AST work-load.

The remaining structure of this paper is as follows: Section 2 provides information about the database used to develop ML algorithms. Section 3 describes the utilized ML-based methodology for the prediction of the presence of genes and pathotypes. Section 4 describes the results achieved by the proposed algorithms. Section 5 discusses the obtained results. Finally, the conclusion of the proposed work is presented in section 6.

## 2. Database used: isolates —materials

### 2.1. Bacterial isolation and identification

In this study, for two years (2018-2020), 341 non-duplicate clinical isolates of *K. pneumoniae* were collected from five university hospitals in Tehran and Qazvin, Iran. The isolates were identified as *K. pneumoniae* using conventional standard biochemical tests. Then, the colony of bacterial isolates were suspended in a Nutrient Broth containing 20% glycerol and stored in a -80°C freezer for further experiments.

### 2.2. hv*Kp* identification

Since most hypervirulent *K. pneumoniae* isolates are tellurite resistant and carry aerobactin genes (*iucA* and *iutA*), we used these properties to identify them, as described in our previous study (14). The detection of *iucA* and *iutA* in hv*Kp* isolates was PCR-based (*iucA* forward, AATCAATGGCTATTCCCGCTG; *iucA* reverse, CGCTTCACTTCTTTCACTGACAGG; *iutA* forward, GCCGCTAGGTTGGTGATGT; *iutA* reverse, CTCTGGTCGTGCTGGTTGA) (14).

### 2.3. Antimicrobial susceptibility test

In this part of the study, the 341 *K. pneumoniae* isolates (including 102 hv*Kp* and 239 c*Kp* isolates) were utilized for antibiotic susceptibility testing. The disk diffusion method was applied to determine the antimicrobial susceptibility testing according to Clinical and Laboratory Standards Institute recommendations (CLSI, 2018) for the 341 *K. pneumoniae* isolates. The antibiotic discs used in this study are as follows: Amikacin/ 30 μg (AK), Gentamicin / 10 μg (GN), Cefotaxime/ 30 μg (CTX), Ceftazidime/ 30 μg (CAZ), Ceftriaxone/ 30 μg (CRO), Imipenem/ 10 μg (IMI), Meropenem/ 10 μg (MEM), Cefepime/ 30 μg (FEP), Ciprofloxacin/ 30 μg (CIP), Ampicillin/ 30 μg (AMP) and Aztreonam/ 30 μg (AZM).

*Escherichia coli* as reference strain ATCC 25922 was used for quality control of antimicrobial disk susceptibility tests. According to the CLSI instructions (2018), the strains that showed non-susceptibility to at least one agent in three classes of antimicrobials were considered MDR. The highest rate of antibiotic resistance was related to ampicillin (100%), followed by cefotaxime (76.2%). The lowest rate of resistance was found in meropenem (24.3%). The resistance rates to other antibiotics including ceftazidime, cefepime, ciprofloxacin, ceftriaxone, gentamicin, imipenem, aztreonam, and amikacin were 75.9%, 62.7%, 59.5%, 54.2%, 52.7%, 44.3%, 39.3%, and 24.6% respectively. In addition, 40.1% of the isolates were resistant to at least three classes of antibiotics and were defined as multidrug-resistant (MDR) (15).

### 2.4. Detection of antimicrobial resistance genes (AMR)

All 341 *K. pneumoniae* isolates were subjected to genotypic identification of beta-lactamase genes using the PCR method by the specific primers (Table 1).

**Table 1.**
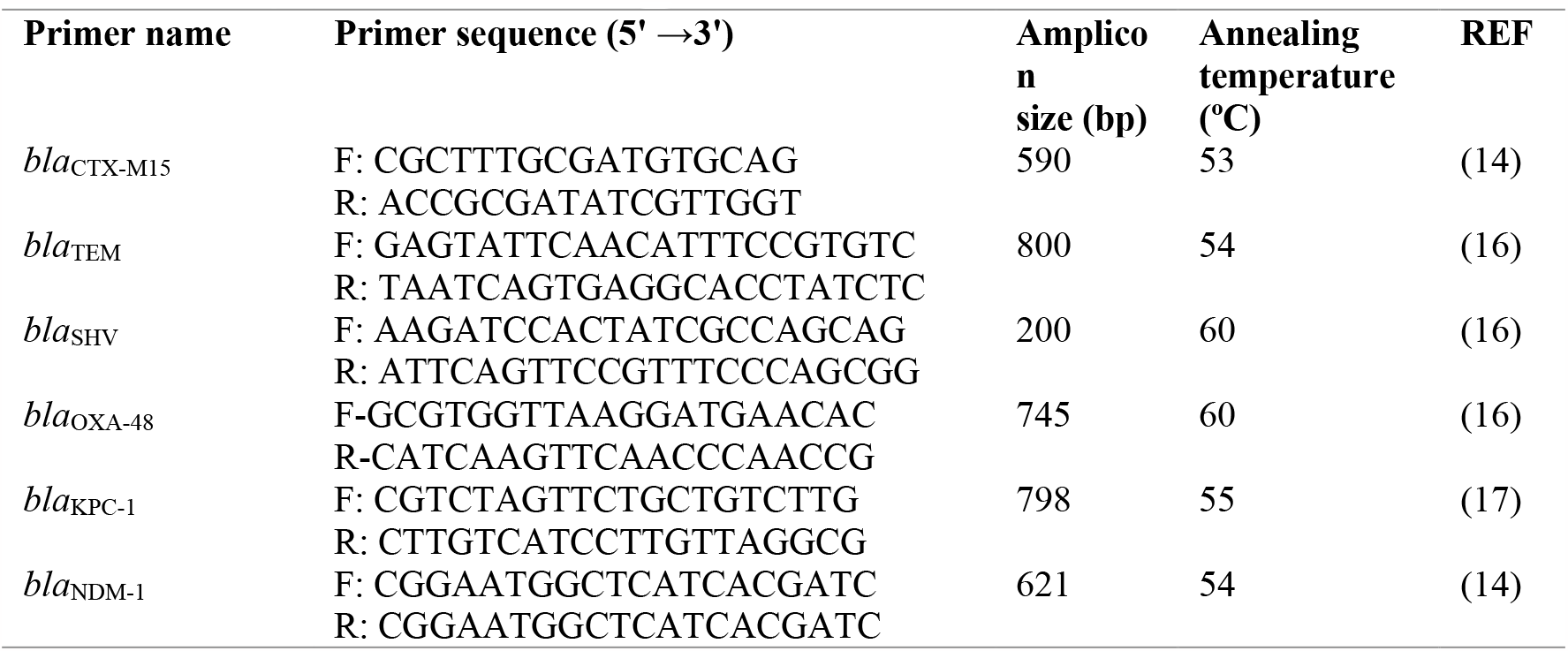
Primer Sequences Used in This Study.

### 2.5. The distribution of data

As mentioned in the previous sections, this study has two goals: detecting *K. pneumonia* pathotypes and detecting the presence of antimicrobial resistance genes. There are two pathotypes of *K. pneumoniae*: hv*Kp* and c*Kp*. Each of the pathotypes can have multiple antibiotic resistance, so we are facing four pathotypes including: multi-drug resistance hypervirulent *K. pneumonia* (MDR-hv*Kp*), non-MDR hv*Kp*, multi-drug resistance c*Kp* (MDR-c*Kp*), and none-MDR. The distribution and the percentage of the pathotype have been demonstrated in Fig. 1. Regarding the second goal of the study, the detection of beta-lactamase genes is assumed. These genes are *bla*_CTX-M15_, *bla*_TEM_, *bla*_SHV_, *bla*_OXA-48_, and *bla*_NDM-1_. The histogram of the genes based on their pathotype classes has been shown in Fig. 2.

**Fig. 1.**
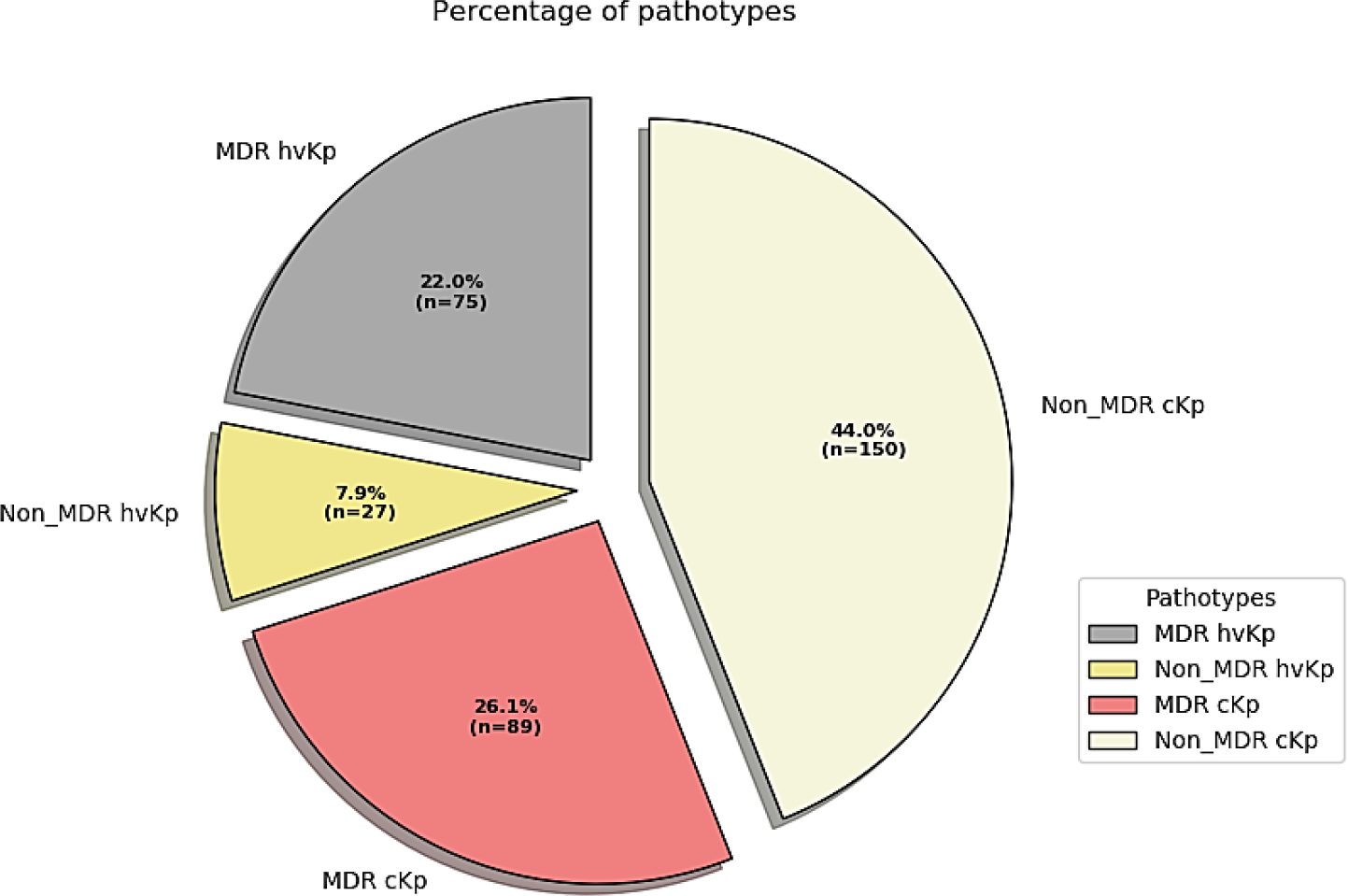
The distribution of Pathotypes

**Fig. 2.**
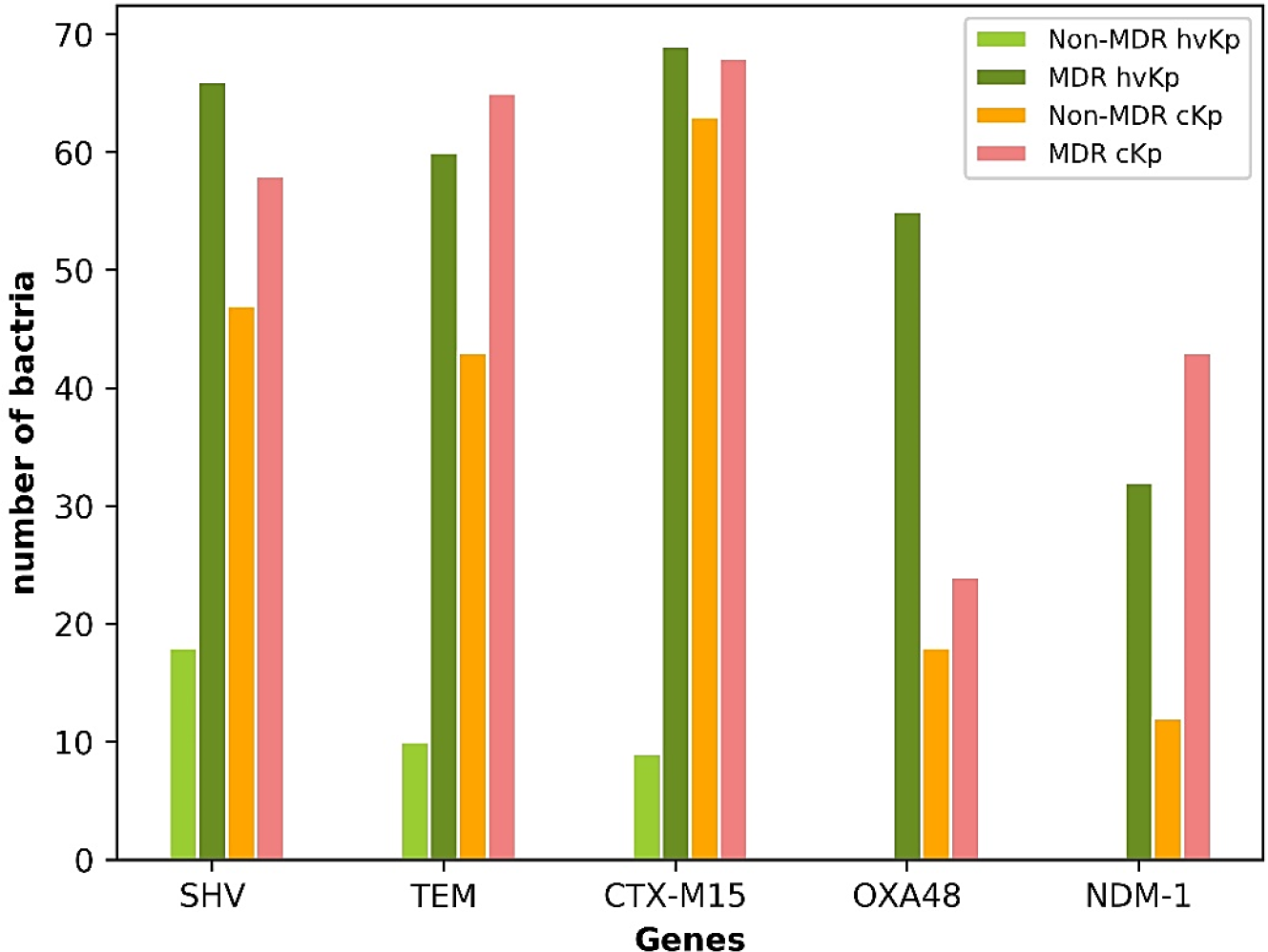
The histogram of antimicrobial resistance genes.

Moreover, a comprehensive histogram for the susceptibility of isolates to all antibiotics has been presented in Fig. 3.

**Fig. 3.**
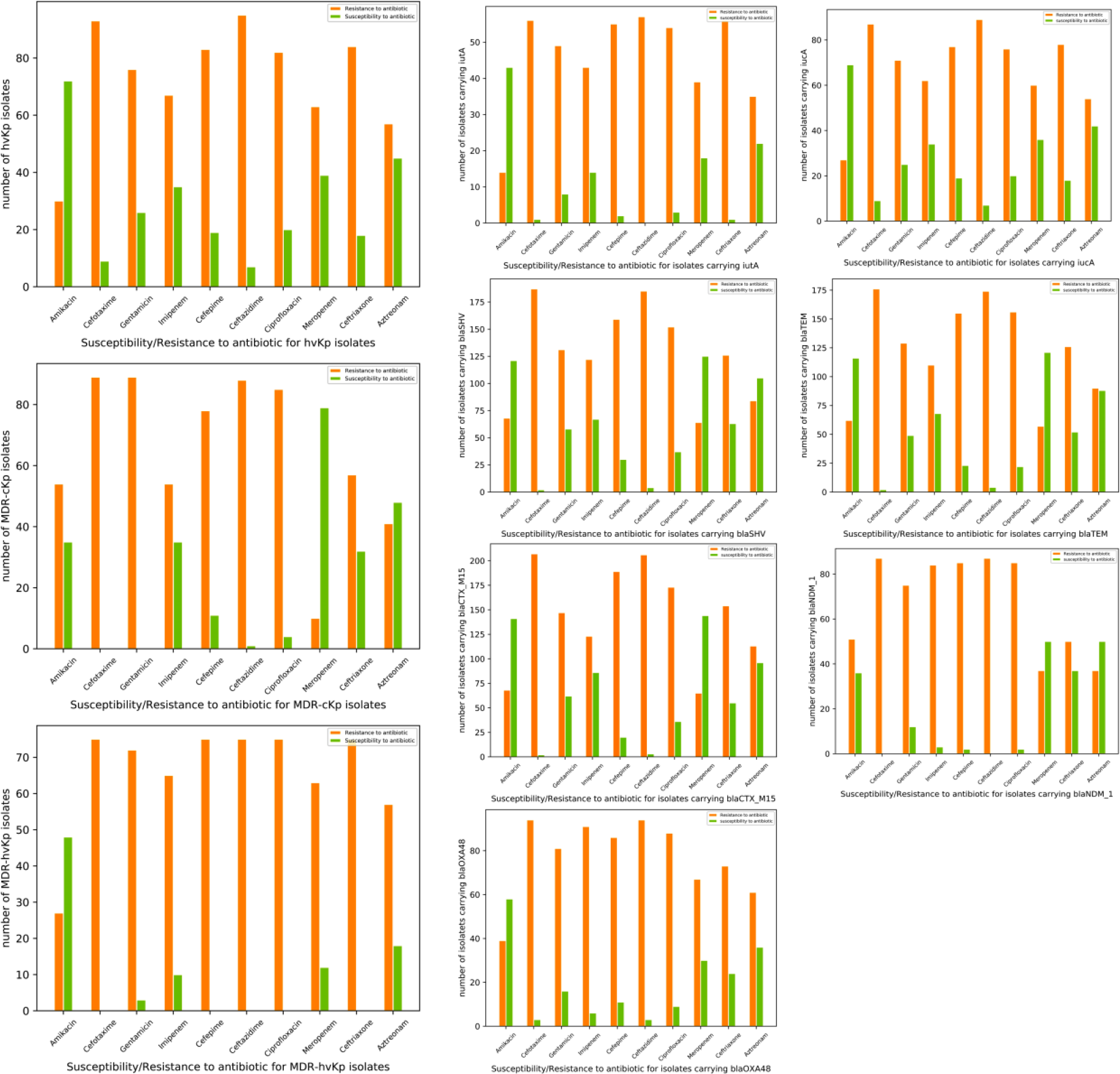
The histogram of resistance/susceptibility of the isolates to all antibiotics

## 3. Overview of the proposed method

Generally, our method consists of four steps. The first step is about data collection and biological experiments in which 341 non-duplicated isolates are used, then the molecular diagnosis of two pathotypes (hv*Kp* and c*Kp*) followed by an antimicrobial susceptibility test and determination of MDR isolates is performed. The second step is preprocessing that mainly contains feature selection algorithms which are Pearson’s Chi-squared test and the Wald test. The third step is to train classification models alongside a model-agnostic approach as a wrapper method for feature selection. There are five ML algorithms selected to be trained on the training dataset including, Logistic Regression (LR), Naïve Bayes Classifier (NBC), Linear Discriminant Analysis (LDA), Support Vector Machine (SVM), and CatBoost. And, the fourth step is to specify crucial antibiotics for the detection of each pathotype or gene by applying training models to test the data set and comparing their performance based on selected antibiotics. Fig. 4 shows the flowchart diagram of the proposed framework.

**Fig. 4.**
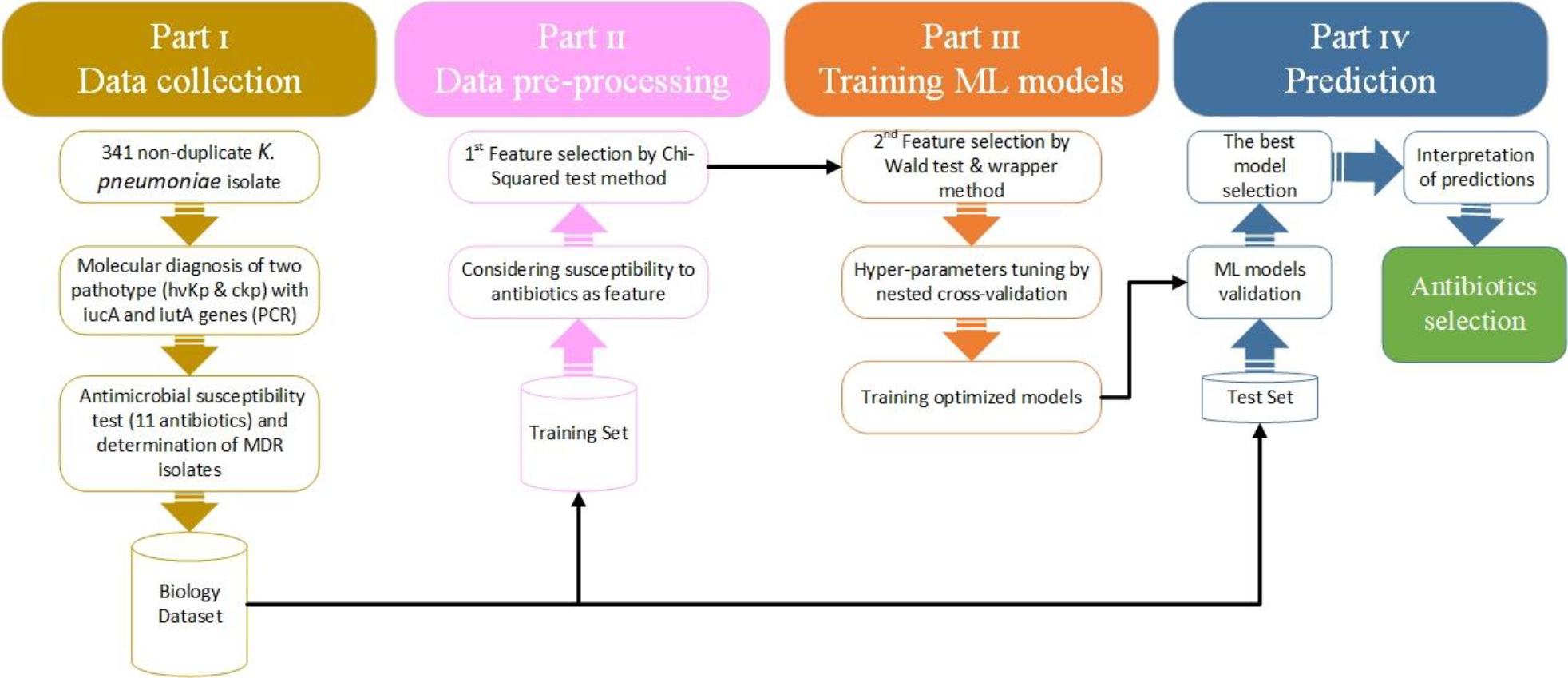
Scheme of machine learning (ML)-based approaches for the gene prediction

It is worth mentioning that the features used as the input of ML models are actually the result of biological experiments (antimicrobial susceptibility test). Therefore, we have not extracted any handcrafted features through mathematical or pattern recognition approaches. We only selected these biological features. Statistical relationships between antibiotics resistance/susceptibility (features) and the presence of genes or pathotypes (response variables) are specified based on Pearson’s Chi-Squared test (18) for primary feature selection. Then, the Wald test (19) and a Wrapper method (20) are applied to the selected features for secondary feature selection. The primary purpose of selecting the discriminatory features is to obtain a more accurate and precise detection of the presence of genes and pathotypes. Pearson’s Chi-Squared test and the Wald test are hypothesis tests (21).

The ML algorithms were implemented on a desktop computer using R language and the ‘MASS’, ‘naivebayes’, ‘e1071’, ‘DALEX’, ‘catboost’, and ‘caret’ libraries, respectively. The desktop computer has a configuration of Intel CoreTM i7-5500U CPU @ 2.1 GHz processor, 16 GB RAM, 64-bit operating system, and 64-based processor.

### 3.1. Discriminatory feature analysis and prominent feature selection

The accuracy, computational efficiency, and complexity of the ML models depend upon the discriminatory features used in the ML algorithms. If all the extracted features are applied to the algorithm, it may not only affect its accuracy but, at the same time, may also give rise to the complexity and affects CPU time for calculation. Thus, feature selection is crucial as it enables the classifier to train faster on the correct subset of features. It may also improve the accuracy of the developed model.

Filter-based methods (22) including Pearson’s Chi-squared test and Wald test are utilized for feature selection. Pearson’s Chi-squared test is a statistical hypothesis test used for a primary feature selection that examines discrimination for all eleven features (antibiotic susceptibility/resistance). The most effective antibiotics (P-value < 0.05) were selected and utilized in the first model in each algorithm. When the P-value is lower, it shows higher discriminatory power among the features. Antibiotics with a P-Value ≥ 0.05 comply with the Null hypothesis (H0), which means no statistical and significant relationship exists between the presence of genes and pathotypes of isolates with resistance/susceptibility profiles. These non-significant antibiotics through the Chi-Squared test were eliminated at the later feature selection methods (Wald test and model agnostic approach). A second model will be trained for all five ML algorithms based on the selected effective antibiotics via the Wald test as the second filter-based method for feature selection, which shows whether the antibiotic is statistically trivial (Null hypothesis) or this assumption is denied through the alternative hypothesis (P-value < 0.05).

In addition to the previous feature selection methods, a model-agnostic approach for feature selection through DALEX package (23), is employed for applying to LR (19), NBC (24), LDA (25), CatBoost (26), and SVM algorithms (27) to specify effective antibiotics. Also, we consider the simplicity principle for applying the simplest antibiotics combinations to other algorithms.

- Model-agnostic approach; the DALEX package suggests a model-agnostic method to evaluate feature importance based on a permutational approach (23). This package can explore, interpret, and compare predictive models through visualization, which leads to deriving useful information from different predictive models regardless of their internal structure.
- The simplicity principle (28); because of the difference among internal structures of mentioned algorithms and using different feature selection methods for selecting effective antibiotics, we may have different antibiotics combinations in each model. Therefore, based on the simplicity principle, the combinations with the minimum number of antibiotics among all algorithms will be considered to apply to other algorithms in which selected antibiotics combinations for these algorithms differ from the simplest ones.

### 3.2. ML Algorithms and Data Split

Based on the resistance or susceptibility of isolates to the antibiotics, several predictive models by the machine learning algorithms will be trained. These algorithms are LR, NBC, LDA, SVM, and CatBoost. Each model was separately trained on each group of features that selected by the feature selection methods. To train and evaluate the ML models, the dataset was split into training and test sets.

Initially, we divided the dataset into two distinguished portions using one of the sampling methods, including simple random sampling, stratified sampling (29), and the “under and over combination sampling” method (30). According to Fig. 5, since response variables, including *bla*_OXA-48_, *bla*_NDM-1_, hv*Kp*, hv*Kp* -MDR, and *iucA*, are imbalanced, stratified sampling will help ensure a balanced representation of these response variables. However, the dataset is highly biased towards the absence of *iutA* and even using stratified sampling is not useful. Therefore, the “under and over combination sampling” method will be applied to split the dataset into train and test sets. Twenty percent of the dataset was allocated to the test dataset for validation. The rest was dedicated to the training dataset for model training and the crossvalidation process. Hyper parameters (if applicable) will be tuned by employing 10-fold crossvalidation. Then, the test data will assess each model’s performance and generalization ability.

**Fig. 5.**
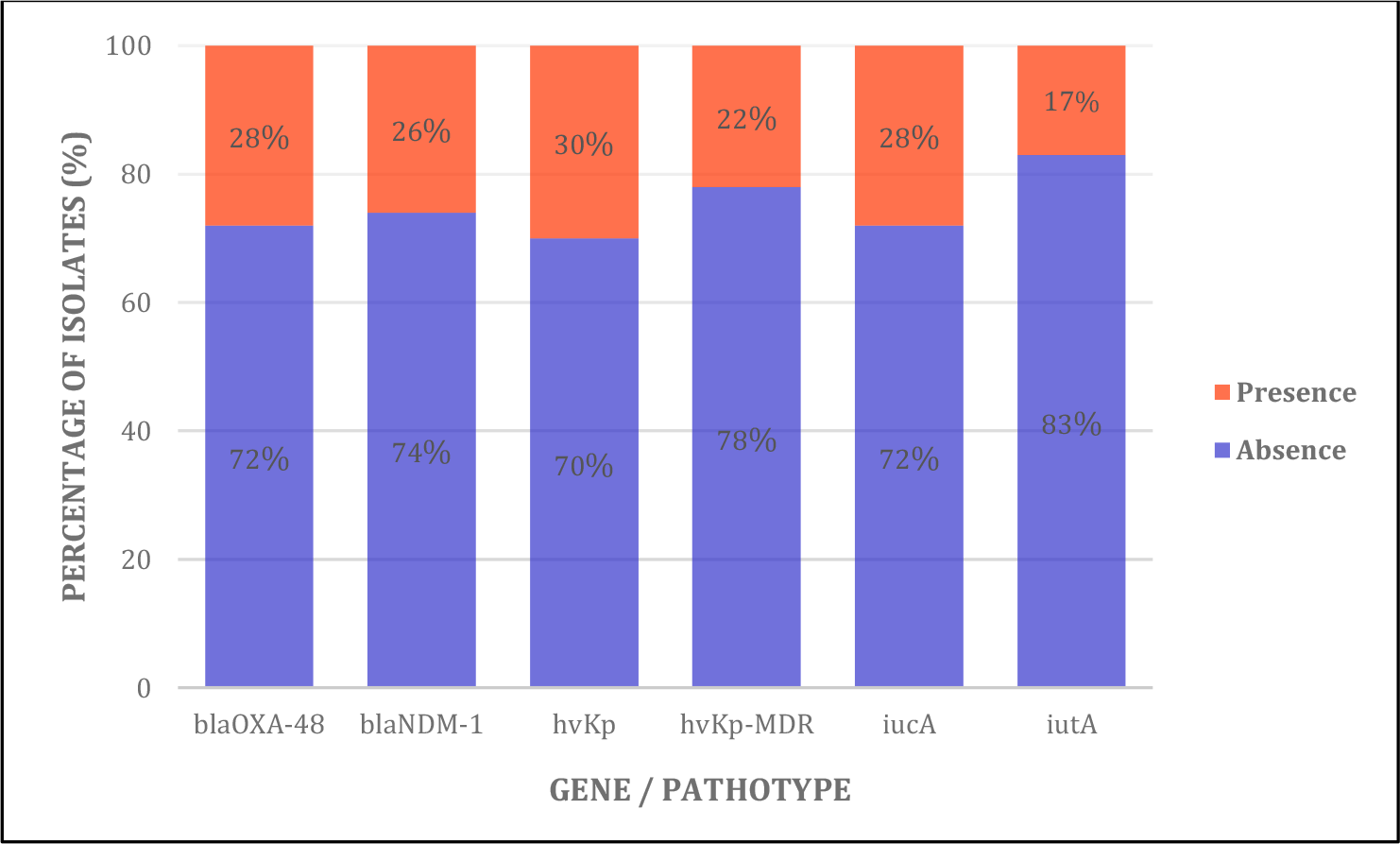
Percentage of isolates in which presence or absence of the specified genes and pathotypes have bias

## 4. Results

### 4.1 Pathotype identification

The differentiation of the two pathotypes c*Kp* and hv*Kp* was made based on the two virulence factor genes including *iucA* and *iutA* which are related to aerobactin as a main siderophore of hypervirulent *K. pneumoniae* isolate. Out of 341 *K*.*pneumoniae* isolates, 102 isolates were identified as hv*Kp* pathotypes. In this way, 46 and 5 isolates carried the *iucA* and *iutA* genes alone respectively, and the simultaneous presence of the *iucA* and *iutA* genes was detected in 51 isolates. The rest of the isolates (n=239), that did not have hv*Kp* diagnostic molecular markers, were reported as c*Kp*.

### 4.2 ML analysis

As mentioned in 2.3, all the isolates in the data set are resistant to Ampicillin, the sole antibiotic with such characterization. Therefore, Ampicillin has not any impact on the prediction process; therefore, it is eliminated from all of the ML algorithms’ input variables.

Tables 2 to 11 represent the performance of models concerning sensivity, precision, and F1-score. In order to take into account models’ uncertainty, all the values are reported with a 95% confidence interval using the Clopper-Pearson method (31). Confidence interval for F1-score for a multi-class classification has been computed by (32). Considering some changes shown in Code availability section, we can compute confidence interval for F1-score in a binary classification.

**Table 2.**
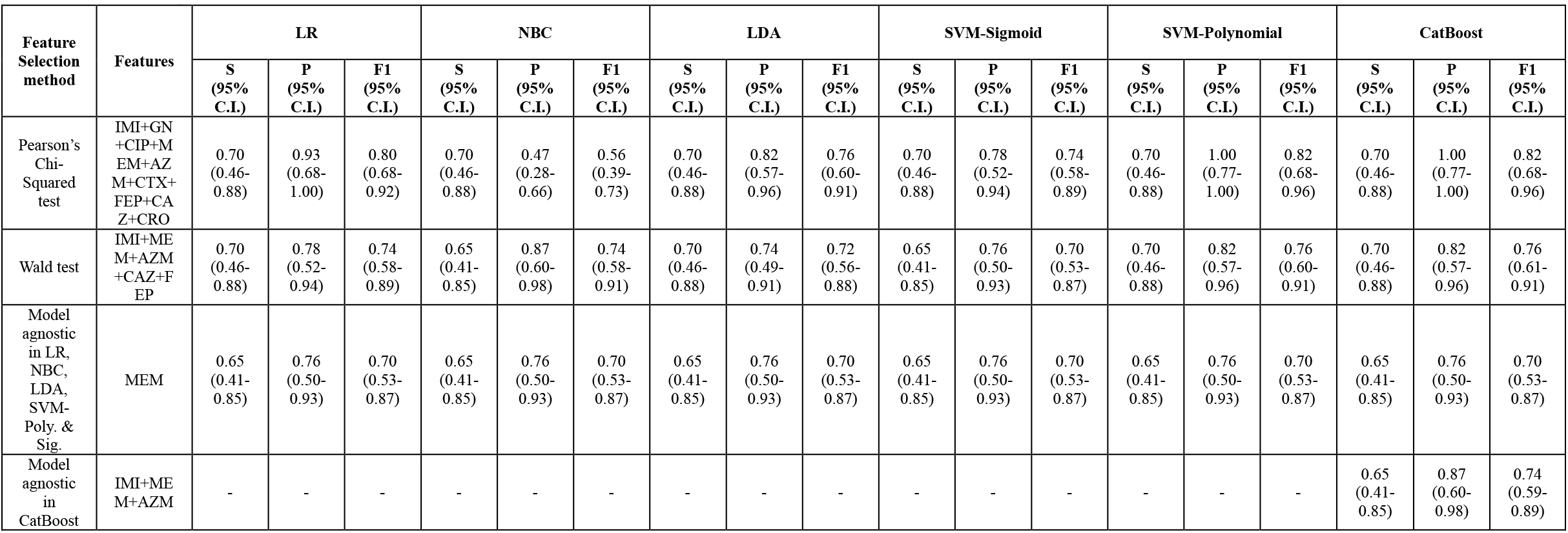
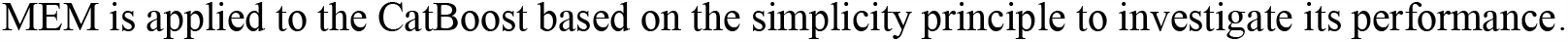
Performance of ML Models predicting the presence of *iucA* in the test dataset based on sensitivity, precision, and F1-Score.

| Feature Selection method | Features | LR |  |  | NBC |  |  | LDA |  |  | SVM-Sigmoid |  |  | SVM-Polynomial |  |  | CatBoost |  |  |
| --- | --- | --- | --- | --- | --- | --- | --- | --- | --- | --- | --- | --- | --- | --- | --- | --- | --- | --- | --- |
|  |  | S (95% C.I.) | P (95% C.I.) | F1 (95% C.I.) | S (95% C.I.) | P (95% C.I.) | F1 (95% C.I.) | S (95% C.I.) | P (95% C.I.) | F1 (95% C.I.) | S (95% C.I.) | P (95% C.I.) | F1 (95% C.I.) | S (95% C.I.) | P (95% C.I.) | F1 (95% C.I.) | S (95% C.I.) | P (95% C.I.) | F1 (95% C.I.) |
| Pearson's Chi-Squared test | IMI+GN+CIP+MEM+AZM+CTX+FEP+CAZ+CRO | 0.70<br>(0.46-0.88) | 0.93<br>(0.68-1.00) | 0.80<br>(0.68-0.92) | 0.70<br>(0.46-0.88) | 0.47<br>(0.28-0.66) | 0.56<br>(0.39-0.73) | 0.70<br>(0.46-0.88) | 0.82<br>(0.57-0.96) | 0.76<br>(0.60-0.91) | 0.70<br>(0.46-0.88) | 0.78<br>(0.52-0.94) | 0.74<br>(0.58-0.89) | 0.70<br>(0.46-0.88) | 1.00<br>(0.77-1.00) | 0.82<br>(0.68-0.96) | 0.70<br>(0.46-0.88) | 1.00<br>(0.77-1.00) | 0.82<br>(0.68-0.96) |
| Wald test | IMI+MEM+AZM+CAZ+FEP | 0.70<br>(0.46-0.88) | 0.78<br>(0.52-0.94) | 0.74<br>(0.58-0.89) | 0.65<br>(0.41-0.85) | 0.87<br>(0.60-0.98) | 0.74<br>(0.58-0.91) | 0.70<br>(0.46-0.88) | 0.74<br>(0.49-0.91) | 0.72<br>(0.56-0.88) | 0.65<br>(0.41-0.85) | 0.76<br>(0.50-0.93) | 0.70<br>(0.53-0.87) | 0.70<br>(0.46-0.88) | 0.82<br>(0.57-0.96) | 0.76<br>(0.60-0.91) | 0.70<br>(0.46-0.88) | 0.82<br>(0.57-0.96) | 0.76<br>(0.61-0.91) |
| Model agnostic in LR, NBC, LDA, SVM-Poly. & Sig. | MEM | 0.65<br>(0.41-0.85) | 0.76<br>(0.50-0.93) | 0.70<br>(0.53-0.87) | 0.65<br>(0.41-0.85) | 0.76<br>(0.50-0.93) | 0.70<br>(0.53-0.87) | 0.65<br>(0.41-0.85) | 0.76<br>(0.50-0.93) | 0.70<br>(0.53-0.87) | 0.65<br>(0.41-0.85) | 0.76<br>(0.50-0.93) | 0.70<br>(0.53-0.87) | 0.65<br>(0.41-0.85) | 0.76<br>(0.50-0.93) | 0.70<br>(0.53-0.87) | 0.65<br>(0.41-0.85) | 0.76<br>(0.50-0.93) | 0.70<br>(0.53-0.87) |
| Model agnostic in CatBoost | IMI+MEM+AZM | - | - | - | - | - | - | - | - | - | - | - | - | - | - | - | 0.65<br>(0.41-0.85) | 0.87<br>(0.60-0.98) | 0.74<br>(0.59-0.89) |

MEM is applied to the CatBoost based on the simplicity principle to investigate its performance.

### 4.3. ML results for aerobactin genes and pathotypes prediction

#### 4.3.1. *IucA* aerobactin *genes*

Considering Table 2, ML models based on selected antibiotics by chi-squared test have more optimum performance than other antibiotic(s) combinations. Coefficients of all the selected antibiotics in logistic function are positive, except Imipenem, Aztreonam, and Cefotaxime. In fact, it can be concluded in isolates susceptible to Aztreonam, Cefotaxime and Imipenem and resistant to 8 other antibiotics, the isolate carries *iucA* with a probability of 46-88% with a 95% confidence interval. However, if we consider simplicity principle alongside model performance, then Meropenem-based models are more optimum. Therefore, considering positive coefficient of MEM in logistic function, if an isolate shows resistance to Meropenem, then the isolate carries *iucA* with a probability of 41-85% with a 95% confidence interval. Feature importance results through model-agnostic approach and Performance of all five models based on the selected antibiotics through Pearson’s Chi-Squared test are shown in Fig. 6 and Fig. 7 respectively.

**Fig. 6.**
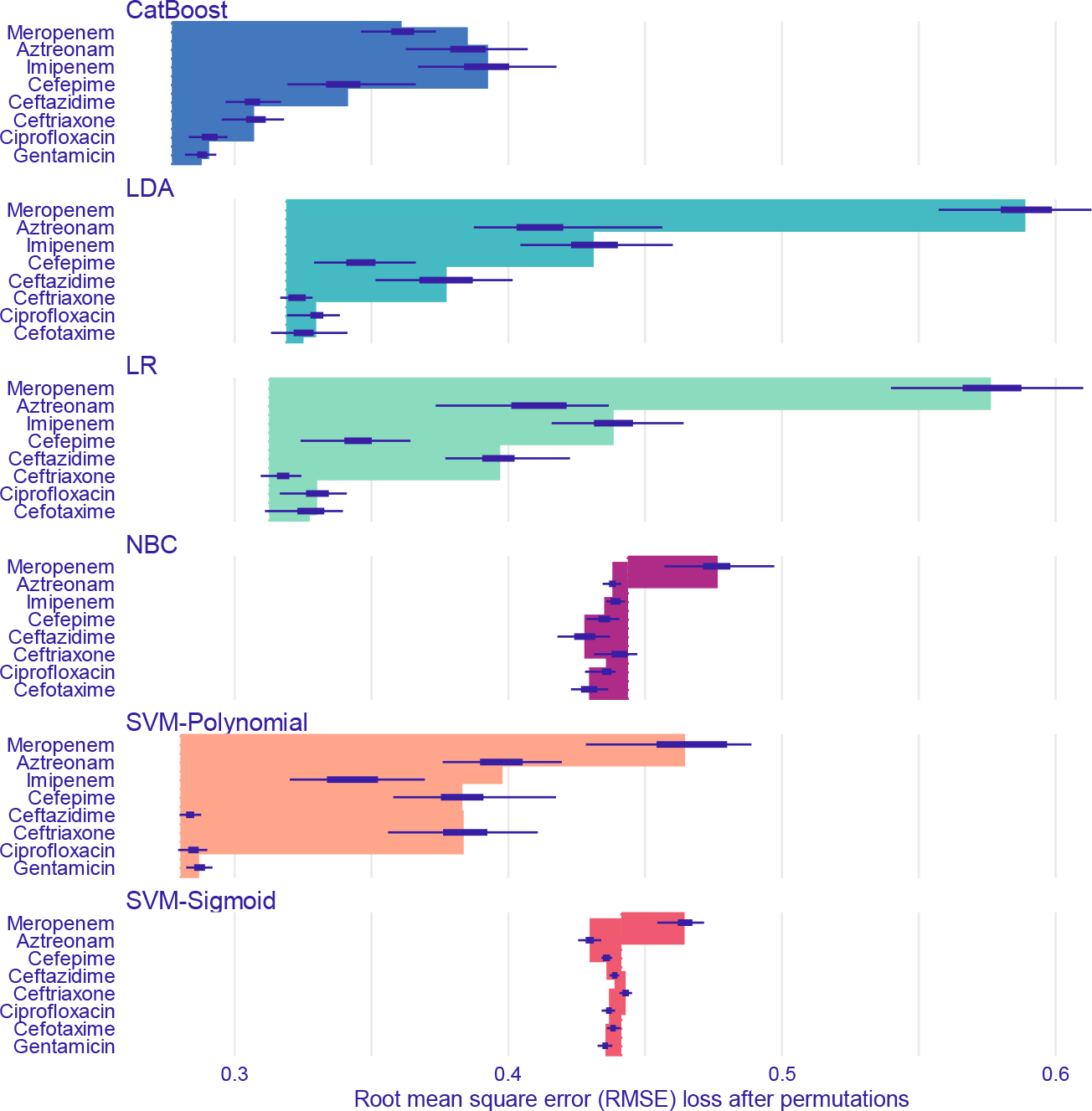
Feature importance of ML models predicting *iucA* aerobactin gene through model-agnostic approach over 50 permutations for top 8 antibiotics

**Fig. 7.**
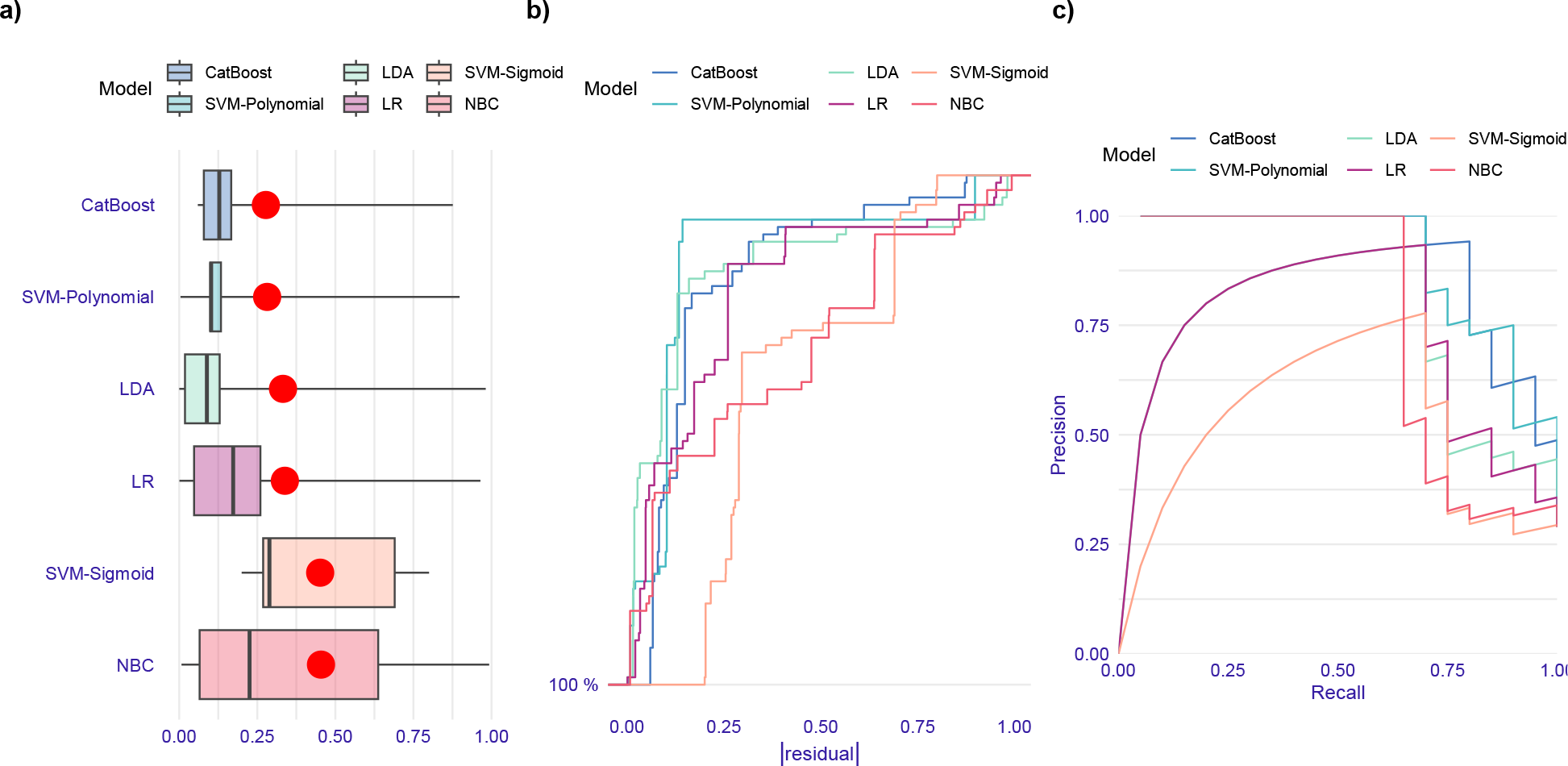
Evaluation of ML models performance in prediction of *iucA* aerobactin gene based on the test data set. (a) Boxplots of absolute values of residuals. Red dot stands for root mean square of residuals. (b) Reverse cumulative distribution of absolute values of residuals. The horizontal axis of the graph represents absolute values of the residuals and the vertical axis represents the percent of isolates having equal to or greater than the specified absolute residual value. Steepness in the plots is a reflection of the variance of residuals and shallower middle sections relative to end sections means larger variance. Moreover, the more a curve tends to downward-sloping straight-line, the more uniform distribution of residuals. CatBoost, LDA, and polynomial SVM models approximately have the same and lowest fraction of the isolates having high residuals. However, sigmoid SVM has the highest fraction of residuals and alongside NBC model have more uniform distribution of residuals than other models. (c) Precision-recall curves. CatBoost and SVM-polynomial models outperform others regarding residual distribution and precision-recall curve.

#### 4.3.2. IutA aerobactin genes

Regarding Table 3, CRO-based, MEM-based models, and the CatBoost model based on the selected antibiotics through the model-agnostic approach including, Ceftriaxone, Aztreonam, Cefepime, and Meropenem, perform better than the other models. Although, performance of MEM-based and CRO-based models in detecting isolates carrying *iutA* (sensitivity values) are close to the CatBoost model, precision value (consequently F1-Score) in the CatBoost model is much higher that causes the CatBoost model is the best model.

**Table 3.** Performance of ML Models predicting the presence of *iutA* in the test dataset based on sensitivity, precision, and F1-Score.

| Feature Selection method | Features | LR |  |  | NBC |  |  | LDA |  |  | SVM-Sigmoid |  |  | SVM-Polynomial |  |  | CatBoost |  |  |
| --- | --- | --- | --- | --- | --- | --- | --- | --- | --- | --- | --- | --- | --- | --- | --- | --- | --- | --- | --- |
|  |  | S (95% C.I.) | P (95% C.I.) | F1 (95% C.I.) | S (95% C.I.) | P (95% C.I.) | F1 (95% C.I.) | S (95% C.I.) | P (95% C.I.) | F1 (95% C.I.) | S (95% C.I.) | P (95% C.I.) | F1 (95% C.I.) | S (95% C.I.) | P (95% C.I.) | F1 (95% C.I.) | S (95% C.I.) | P (95% C.I.) | F1 (95% C.I.) |
| Pearson's Chi-Squared test | IMI+GN+CIP+MEM+AZM+CTX+FEP+CAZ+CRO | 1.00 (0.72-1.00) | 0.61 (0.36-0.83) | 0.76 (0.59-0.93) | 1.00 (0.72-1.00) | 0.28 (0.15-0.45) | 0.44 (0.27-0.61) | 1.00 (0.72-1.00) | 0.58 (0.33-0.80) | 0.73 (0.56-0.91) | 1.00 (0.72-1.00) | 0.35 (0.19-0.55) | 0.52 (0.34-0.71) | 1.00 (0.72-1.00) | 0.69 (0.41-0.89) | 0.81 (0.66-0.97) | 1.00 (0.72-1.00) | 0.58 (0.33-0.80) | 0.73 (0.56-0.91) |
| Wald test | MEM+AZM+CTX+FEP+CRO | 1.00 (0.72-1.00) | 0.58 (0.33-0.80) | 0.73 (0.56-0.91) | 1.00 (0.72-1.00) | 0.38 (0.21-0.58) | 0.55 (0.36-0.74) | 1.00 (0.72-1.00) | 0.52 (0.30-0.74) | 0.69 (0.50-0.87) | 1.00 (0.72-1.00) | 0.58 (0.33-0.80) | 0.73 (0.56-0.91) | 1.00 (0.72-1.00) | 0.65 (0.38-0.86) | 0.79 (0.62-0.95) | 1.00 (0.72-1.00) | 0.65 (0.38-0.86) | 0.79 (0.62-0.95) |
| Model agnostic in LR, LDA, SVM-Poly. & Sig. | MEM | 0.91 (0.59-1.00) | 0.50 (0.27-0.73) | 0.65 (0.45-0.84) | 0.91 (0.59-1.00) | 0.50 (0.27-0.73) | 0.65 (0.45-0.84) | 0.91 (0.59-1.00) | 0.50 (0.27-0.73) | 0.65 (0.45-0.84) | 0.91 (0.59-1.00) | 0.50 (0.27-0.73) | 0.65 (0.45-0.84) | 0.91 (0.59-1.00) | 0.50 (0.27-0.73) | 0.65 (0.45-0.84) | 0.91 (0.59-1.00) | 0.50 (0.27-0.73) | 0.65 (0.45-0.84) |
| Model agnostic in NBC | CRO | 1.00<br>(0.72-1.00) | 0.31<br>(0.16-0.48) | 0.47<br>(0.29-0.64) | 1.00<br>(0.72-1.00) | 0.31<br>(0.16-0.48) | 0.47<br>(0.29-0.64) | 1.00<br>(0.72-1.00) | 0.31<br>(0.16-0.48) | 0.47<br>(0.29-0.64) | 1.00<br>(0.72-1.00) | 0.31<br>(0.16-0.48) | 0.47<br>(0.29-0.64) | 1.00<br>(0.72-1.00) | 0.31<br>(0.16-0.48) | 0.47<br>(0.29-0.64) | 1.00<br>(0.72-1.00) | 0.31<br>(0.16-0.48) | 0.47<br>(0.29-0.64) |
| Model agnostic in CatBoost | CRO+AZ<br>M+FEP+<br>MEM | - | - | - | - | - | - | - | - | - | - | - | - | - | - | - | 1.00<br>(0.72-1.00) | 0.65<br>(0.38-0.86) | 0.79<br>(0.62-0.95) |

MEM and CRO are applied to all five algorithms separately based on the simplicity principle to investigate their performance.

**Table 4.**
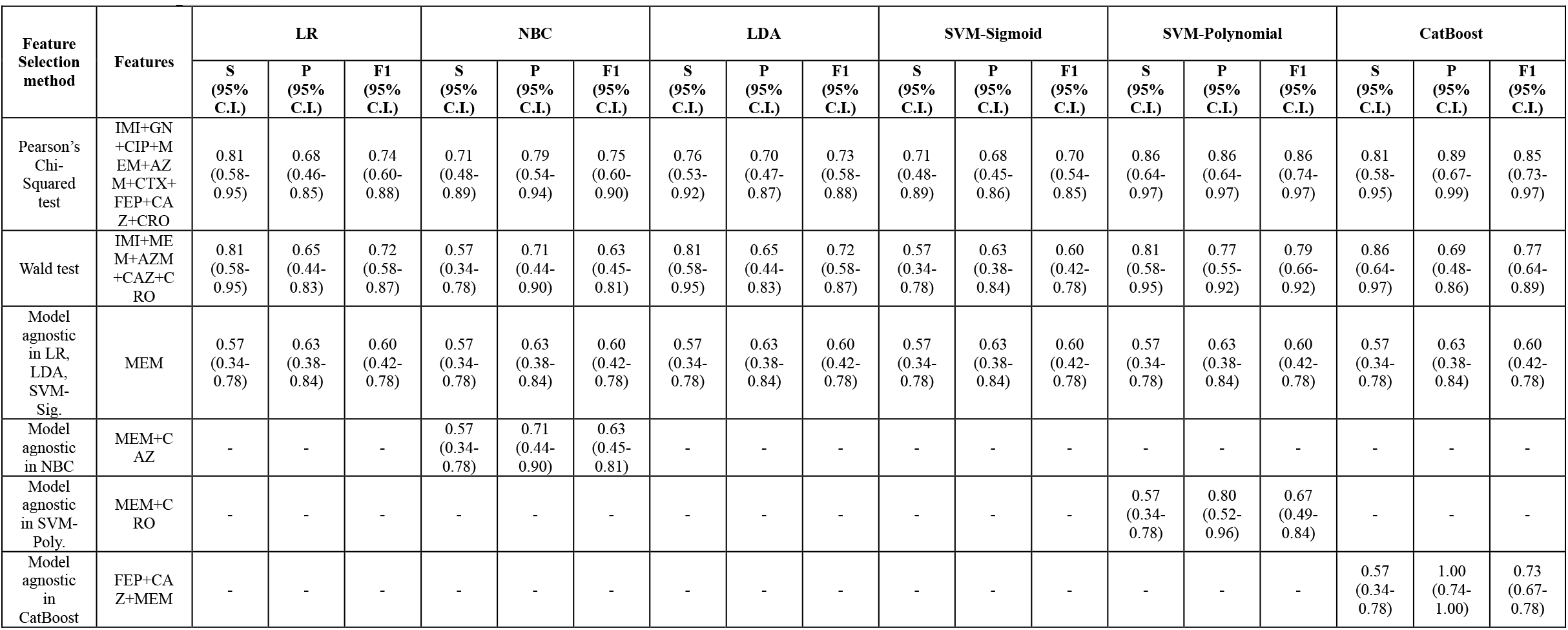
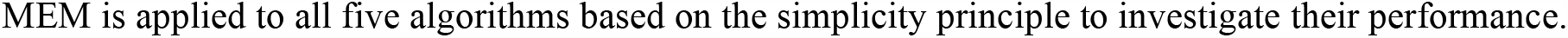
Performance of ML Models predicting the presence of hv*Kp* in the test dataset based on sensitivity, precision, and F1-Score.

Considering the CatBoost mode, due to the negative coefficient of Aztreonam and positive coefficients of Ceftriaxone, Cefepime, and Meropenem in logistic function, if an isolate is susceptible to Aztreonam and resistant to Meropenem, Ceftriaxone, and Cefepime, the isolate carries *iutA* with a probability of 72-100% with a 95% confidence interval that is same as the probability of presence of *iutA* in CRO-based models (sensitivity value). Thus, if detection of isolates carrying *iutA* is the only purpose, only specifying resistance of isolates to Ceftriaxone is required. It should be noted that by considering ceftriaxone solely in the detection, more than half of isolates not carrying *iutA*, classified as the isolates carrying *iutA* (precison).

The results of model-agnostic approach in all algorithms are shown in Fig. 8. Performance of all models considering antibiotics selected by Pearson’s Chi-Squared test are shown in Fig. 9.

**Fig. 8.**
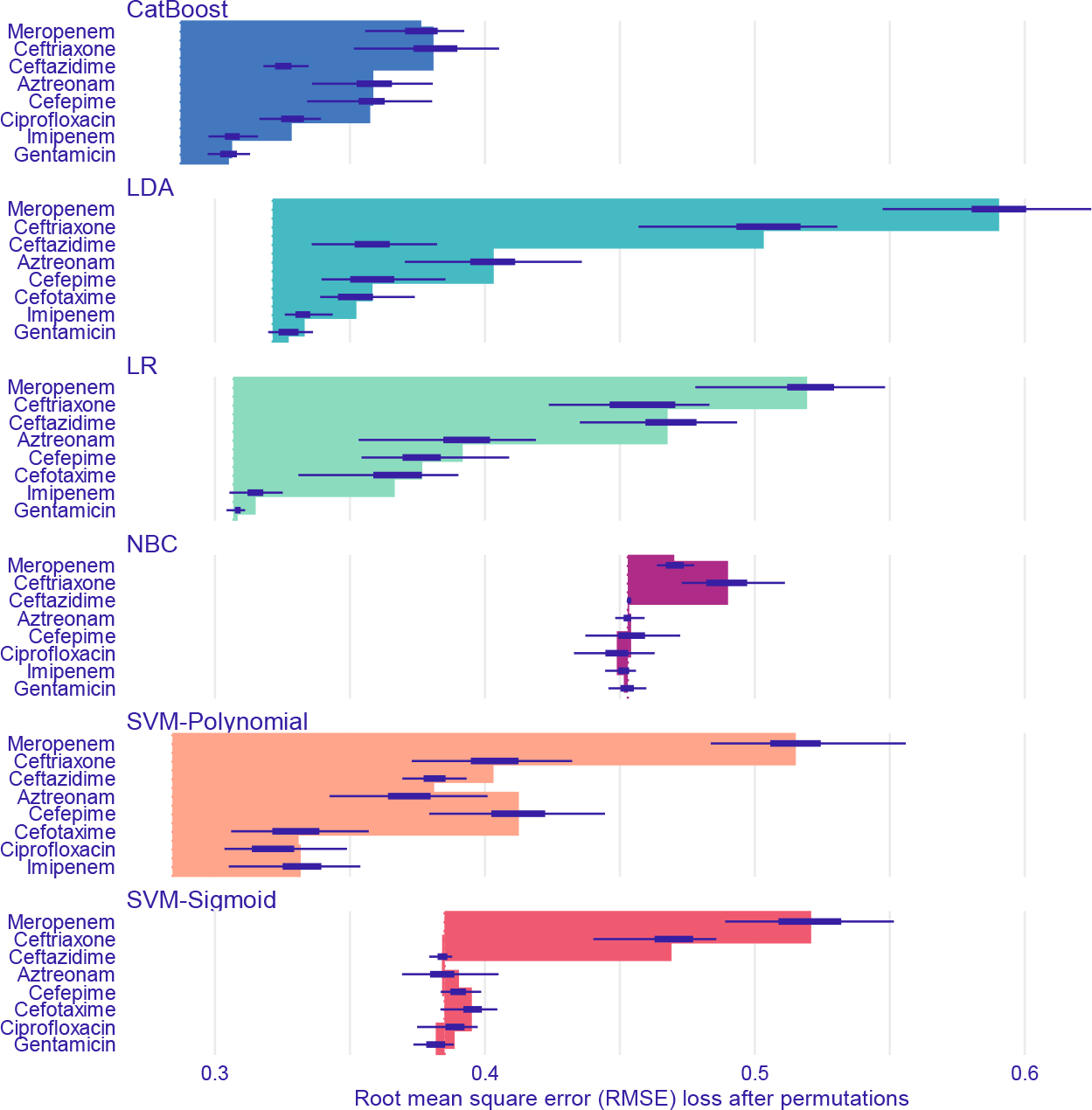
Feature importance of ML models predicting *iutA* aerobactin gene through model-agnostic approach over 50 permutations for top 8 antibiotics

**Fig. 9.**
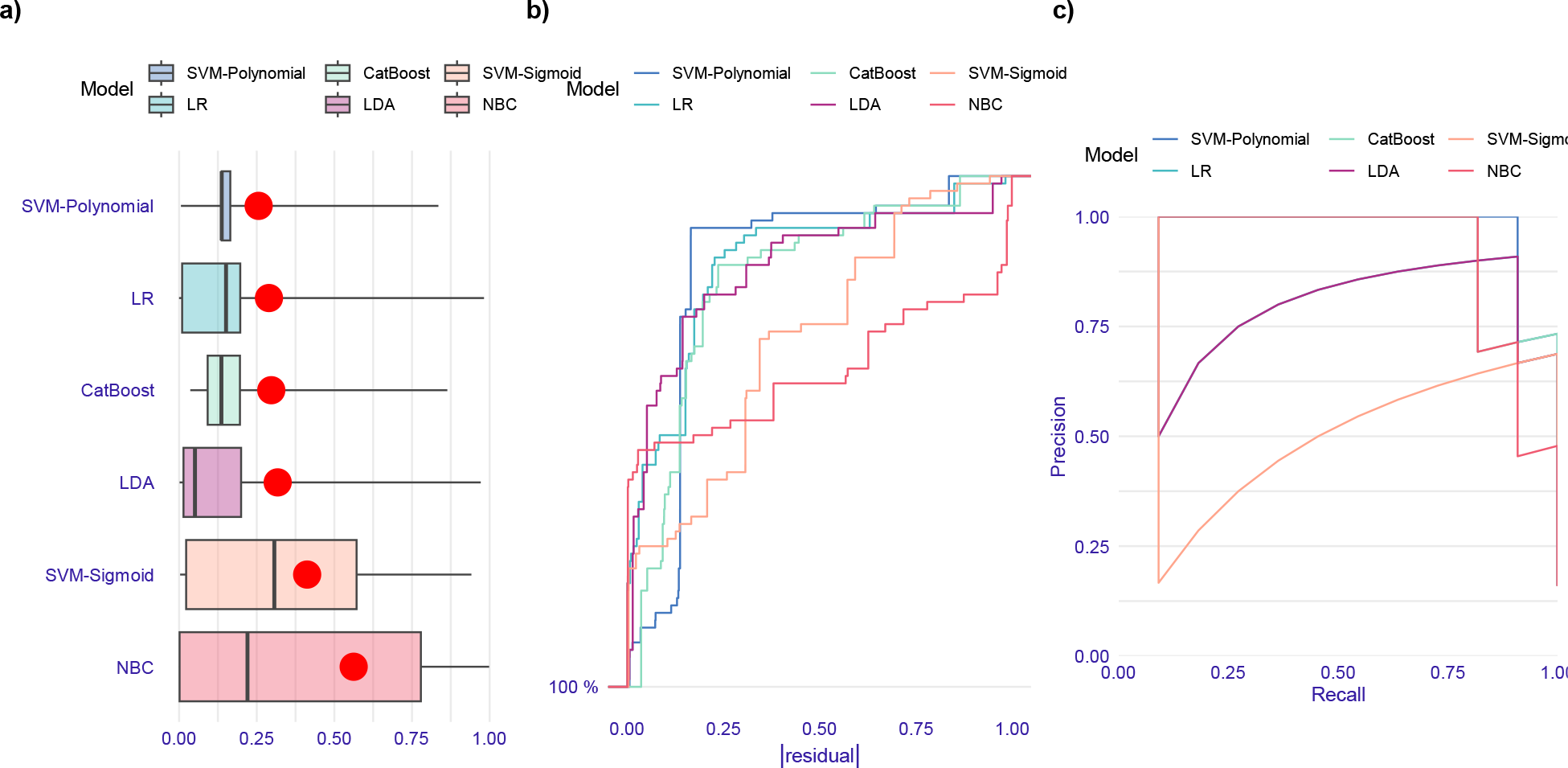
Evaluation of ML models performance in prediction of *iutA* aerobactin gene based on the test data set. (a) Boxplots of absolute values of residuals. Red dot stands for root mean square of residuals. (b) Reverse cumulative distribution of absolute values of residuals. Polynomial SVM model has the lowest fraction of isolates having high residuals and NBC model has the highest fraction. Distribution of residuals in sigmoid SVM model is more uniform than other models. (c) Precision-recall curves. Polynomial kernel of SVM outperforms other models in prediction of iutA gene regarding residual distribution and precision-recall curve.

#### 4.3.3. hvKp pathotype

According to Table 8, polynomial kernel of SVM and CatBoost models based on selected antibiotics through the Wald test perform best. Imipenem and Aztreonam have negative coefficients, but, Meropenem, Ceftazidime, and Ceftriaxone have positive coefficients in logistic function; therefore, if an isolate is susceptible to Imipenem and Aztreonam and shows resistance to Meropenem, Ceftazidime, and Ceftriaxone, the isolate is a hv*Kp* strain with a probability of 64-97% with a 95% confidence interval based on sensitivity value in the CatBoost model.

Crucial antibiotics based on the model-agnostic approach are shown in Fig. 10. Residual distributions and precision-recall curves of all the algorithms based on all the antibiotics except Amikacin are shown in Fig. 11.

**Fig. 10.**
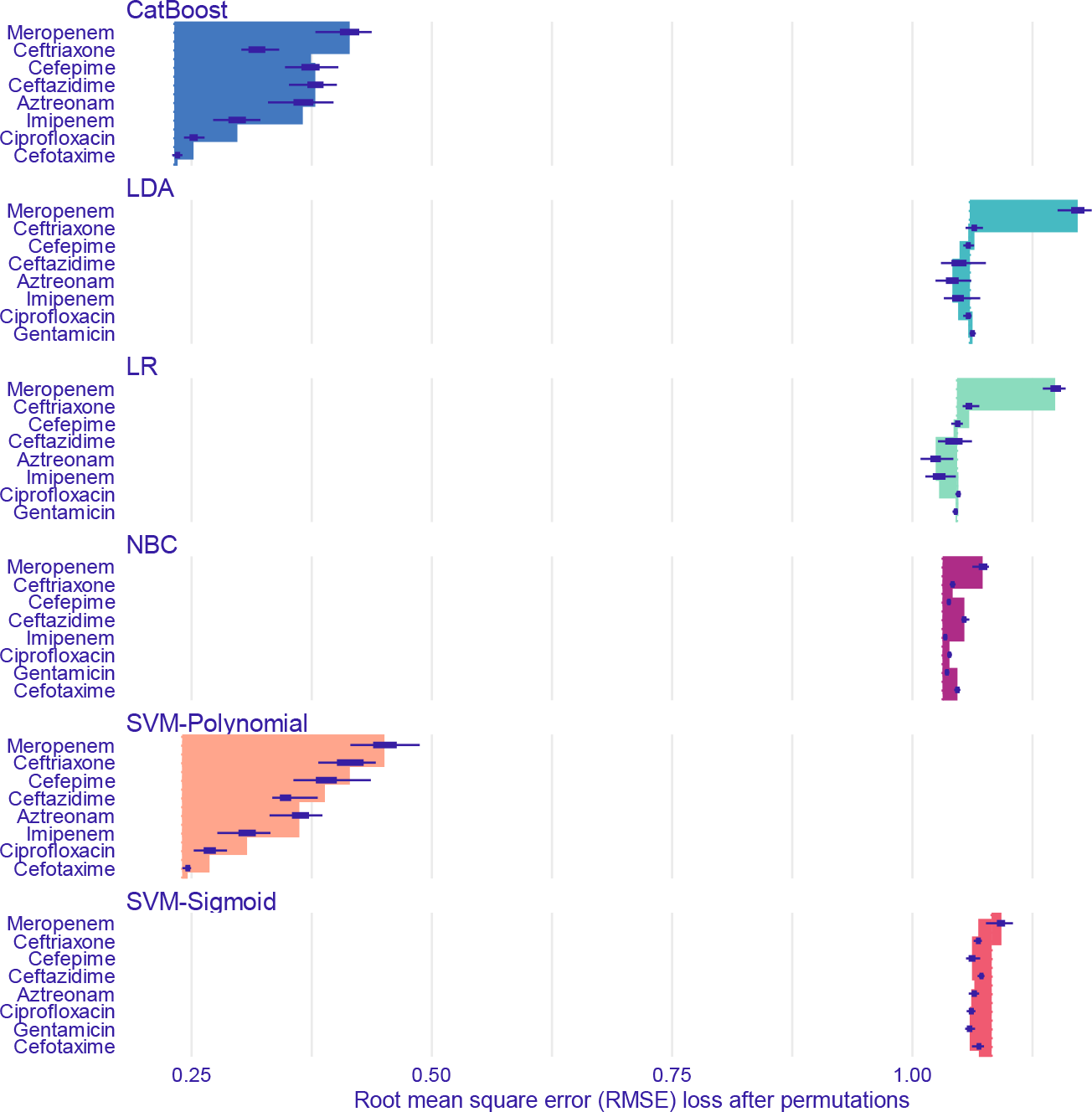
Feature importance of ML models predicting *hvKp* pathotype through model-agnostic approach over 50 permutations for top 8 antibiotics

**Fig. 11.**
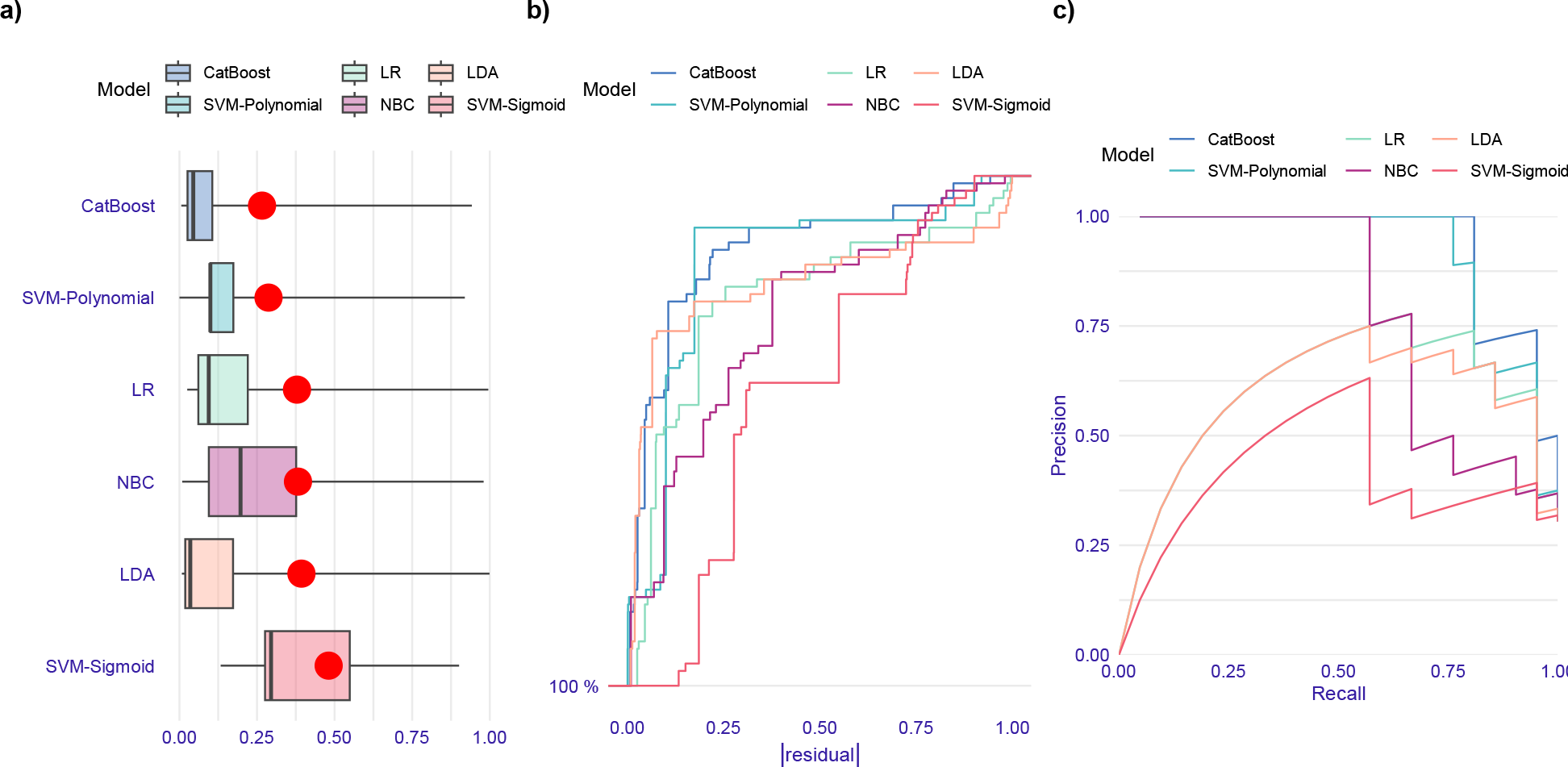
Evaluation of ML models performance in prediction of *hvKp* pathotype based on the test data set. (a) Boxplots of absolute values of residuals. Red dot stands for root mean square of residuals. (b) Reverse cumulative distribution of absolute values of residuals. CatBoost model outperforms other models due to the lowest fraction of the isolates having high residuals. On the contrary, sigmoid SVM has the highest fraction of isolates and this model has more uniform distribution of residuals than other models. (c) Precision-recall curves. CatBoost and SVM-Polynomial models outperform other models

An isolate is a hv*Kp* strain if it carries *iutA* or *iucA* aerobactin genes. We may expect crucial antibiotics for the prediction of hv*Kp* to be comprised of crucial ones for the prediction of *iutA* and *iucA*. Hence, comparing effective antibiotics for the prediction of hv*Kp* to effective antibiotics for the prediction of *iutA* and *iucA* is necessary.

Based on developed ML models, effective antibiotics in detection of hv*Kp* overlap with significant ones in *iutA* and *iucA* genes, including Imipenem, Meropenem, Aztreonam, Ceftriaxone, and Ceftazidime. However, Cefepime as an effective antibiotic for detecting *iucA* and *iutA* genes is not influential in detecting the hv*Kp* pathotype. Thus, the privilege of the ML approach is that we can detect hv*Kp* directly by considering the resistance/susceptibility of an isolate to definite antibiotics that are more limited than effective antibiotics in *iutA* and *iucA*.

#### 4.3.4. MDR-cKp pathotype

According to Table 5, the most optimized model considering quantities of sensitivity, precision, and F1-Score and the simplicity principle is the LR model constructed based on Gentamicin, Ciprofloxacin, Amikacin, Meropenem, and Cefotaxime. The coefficients of these antibiotics in logistic function are positive, thus, isolate’s resistance to these antibiotics results in the presence of MDR-c*Kp* with a probability of 89-100% with a 95% confidence interval. There is a negligible difference between this model and Gentamicin-based models. The sensitivity value for Gentamicin-based models is 1; therefore, the resistance of an isolate to Gentamicin results in the presence of MDR-c*Kp* strain with a probability of with a 95% confidence interval. We can conclude that isolate’s resistance to Gentamicin is sufficient for detection of MDR-c*Kp* strains.

**Table 5.**
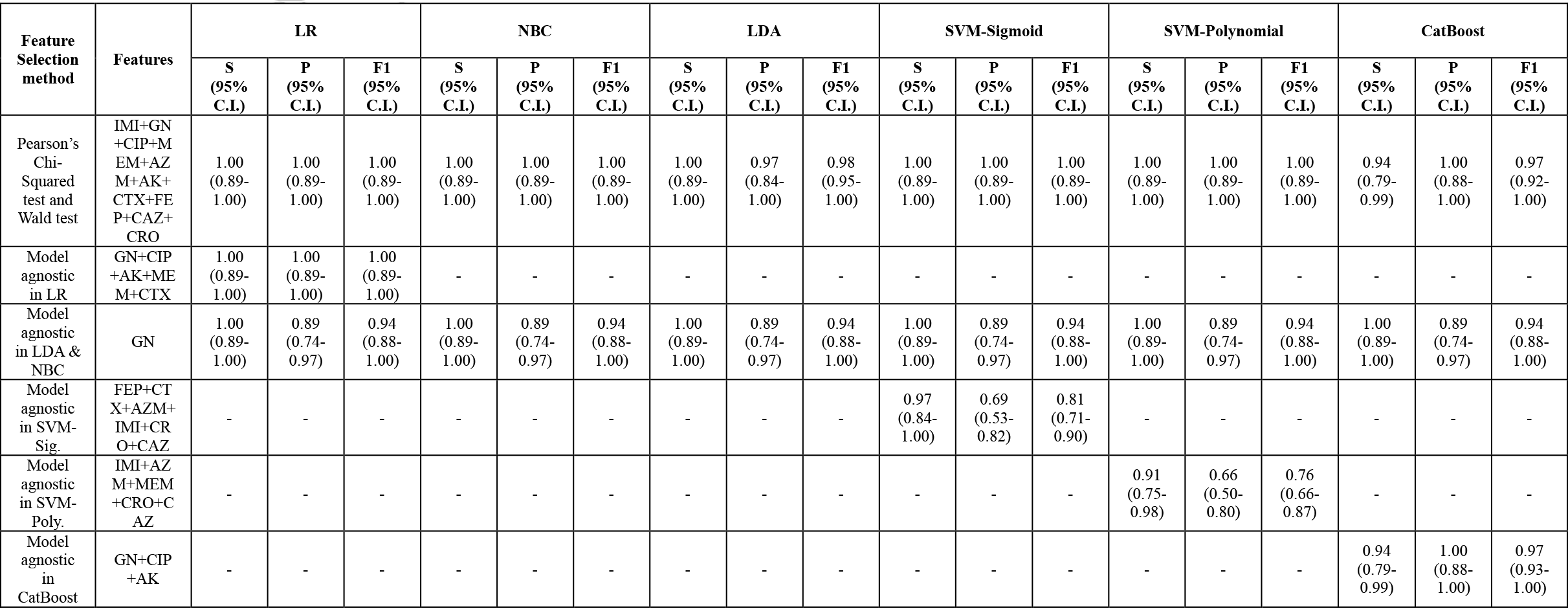
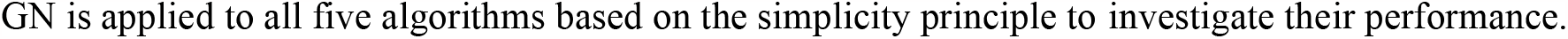
Performance of ML Models predicting the presence of *MDR-cKp* in the test dataset based on sensitivity, precision, and F1-Score.

| Feature Selection method | Features | LR |  |  | NBC |  |  | LDA |  |  | SVM-Sigmoid |  |  | SVM-Polynomial |  |  | CatBoost |  |  |
| --- | --- | --- | --- | --- | --- | --- | --- | --- | --- | --- | --- | --- | --- | --- | --- | --- | --- | --- | --- |
|  |  | S (95% C.I.) | P (95% C.I.) | F1 (95% C.I.) | S (95% C.I.) | P (95% C.I.) | F1 (95% C.I.) | S (95% C.I.) | P (95% C.I.) | F1 (95% C.I.) | S (95% C.I.) | P (95% C.I.) | F1 (95% C.I.) | S (95% C.I.) | P (95% C.I.) | F1 (95% C.I.) | S (95% C.I.) | P (95% C.I.) | F1 (95% C.I.) |
| Pearson's Chi-Squared test and Wald test | IMI+GN+CIP+M<br>EM+AZ<br>M+AK+<br>CTX+FE<br>P+CAZ+<br>CRO | 1.00<br>(0.89-<br>1.00) | 1.00<br>(0.89-<br>1.00) | 1.00<br>(0.89-<br>1.00) | 1.00<br>(0.89-<br>1.00) | 1.00<br>(0.89-<br>1.00) | 1.00<br>(0.89-<br>1.00) | 1.00<br>(0.89-<br>1.00) | 0.97<br>(0.84-<br>1.00) | 0.98<br>(0.95-<br>1.00) | 1.00<br>(0.89-<br>1.00) | 1.00<br>(0.89-<br>1.00) | 1.00<br>(0.89-<br>1.00) | 1.00<br>(0.89-<br>1.00) | 1.00<br>(0.89-<br>1.00) | 1.00<br>(0.89-<br>1.00) | 0.94<br>(0.79-<br>0.99) | 1.00<br>(0.88-<br>1.00) | 0.97<br>(0.92-<br>1.00) |
| Model agnostic in LR | GN+CIP<br>+AK+ME<br>M+CTX | 1.00<br>(0.89-1.00) | 1.00<br>(0.89-1.00) | 1.00<br>(0.89-1.00) | - | - | - | - | - | - | - | - | - | - | - | - | - | - | - |
| Model agnostic in LDA & NBC | GN | 1.00<br>(0.89-1.00) | 0.89<br>(0.74-0.97) | 0.94<br>(0.88-1.00) | 1.00<br>(0.89-1.00) | 0.89<br>(0.74-0.97) | 0.94<br>(0.88-1.00) | 1.00<br>(0.89-1.00) | 0.89<br>(0.74-0.97) | 0.94<br>(0.88-1.00) | 1.00<br>(0.89-1.00) | 0.89<br>(0.74-0.97) | 0.94<br>(0.88-1.00) | 1.00<br>(0.89-1.00) | 0.89<br>(0.74-0.97) | 0.94<br>(0.88-1.00) | 1.00<br>(0.89-1.00) | 0.89<br>(0.74-0.97) | 0.94<br>(0.88-1.00) |
| Model agnostic in SVM-Sig. | FEP+CT<br>X+AZM+<br>IMI+CR<br>O+CAZ | - | - | - | - | - | - | - | - | - | 0.97<br>(0.84-<br>1.00) | 0.69<br>(0.53-<br>0.82) | 0.81<br>(0.71-<br>0.90) | - | - | - | - | - | - |
| Model agnostic in SVM-Poly. | IMI+AZ<br>M+MEM<br>+CRO+C<br>AZ | - | - | - | - | - | - | - | - | - | - | - | - | 0.91<br>(0.75-0.98) | 0.66<br>(0.50-0.80) | 0.76<br>(0.66-0.87) | - | - | - |
| Model agnostic in CatBoost | GN+CIP<br>+AK | - | - | - | - | - | - | - | - | - | - | - | - | - | - | - | 0.94<br>(0.79-0.99) | 1.00<br>(0.88-1.00) | 0.97<br>(0.93-1.00) |

GN is applied to all five algorithms based on the simplicity principle to investigate their performance.

The most important antibiotics based on model-agnostic approach are shown in Fig. 12. Performance of all five models based on the first combination (Pearson’s Chi-Squared test results) are shown in Fig. 13 and

**Fig. 12.**
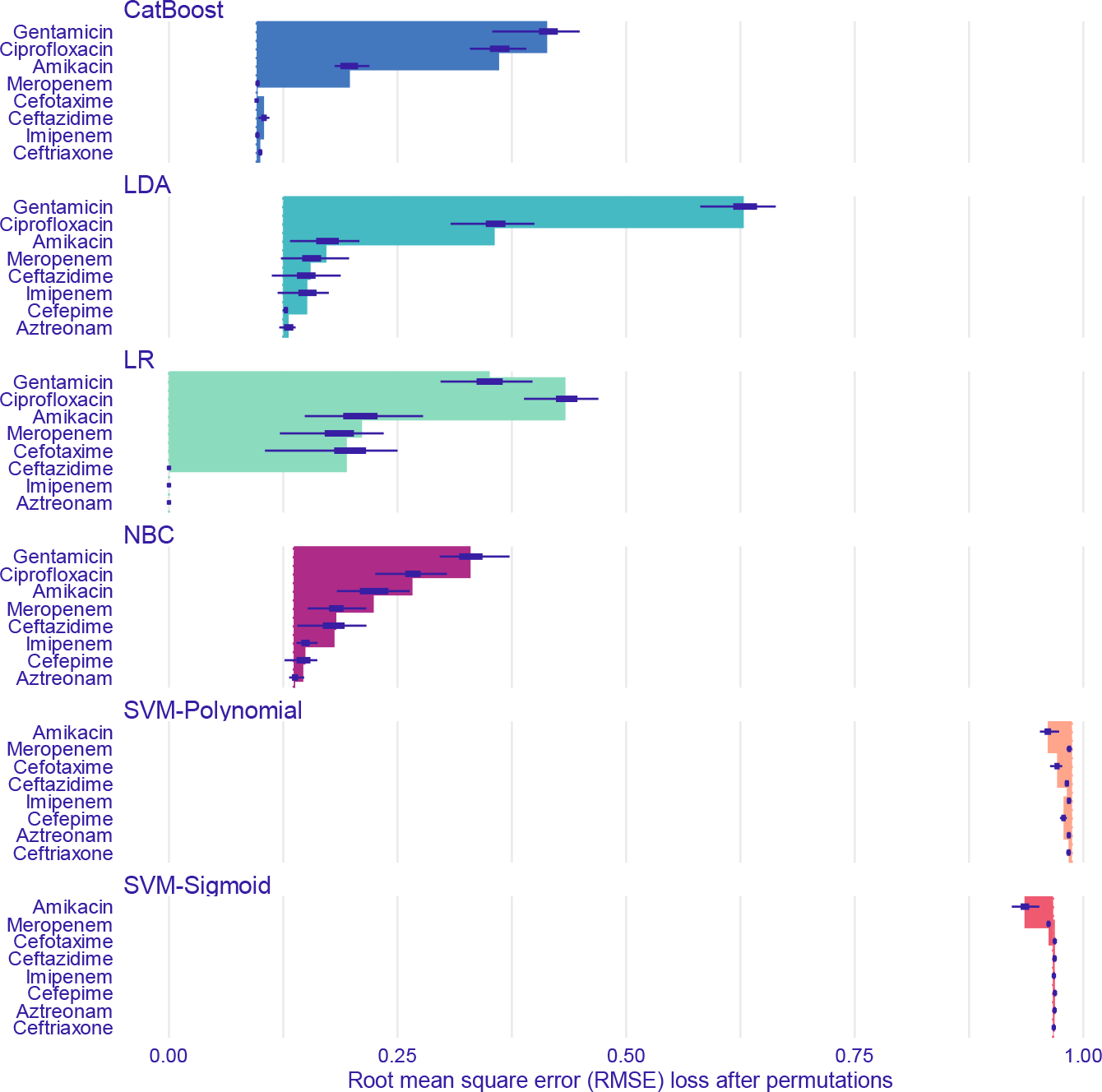
Feature importance of ML models predicting *MDR-cKp* pathotype through model-agnostic approach over 50 permutations for top 8 antibiotics

**Fig. 13.**
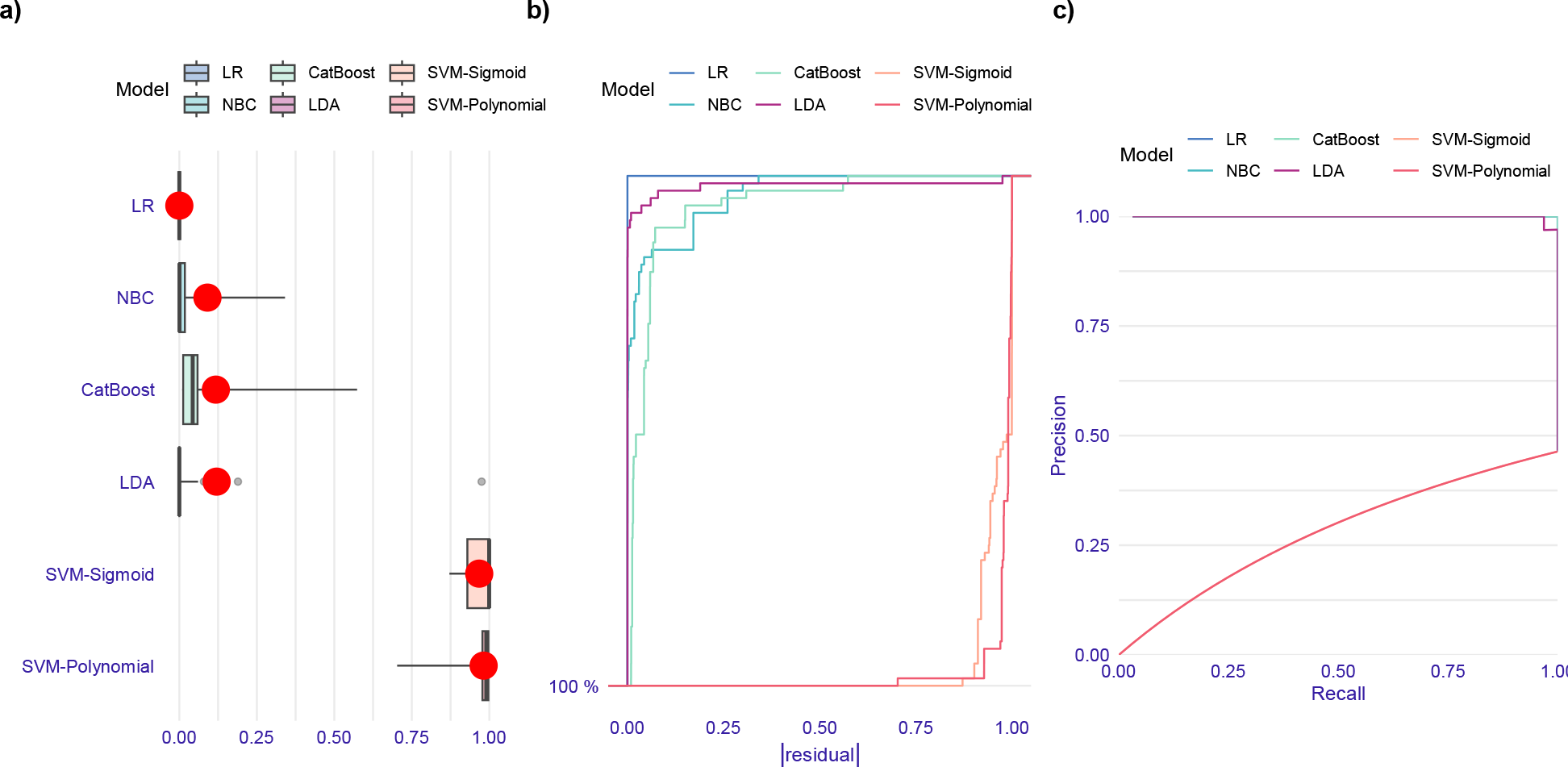
Evaluation of ML models performance in prediction of *MDR-cKp* pathotype based on the test data set. (a) Boxplots of absolute values of residuals. Red dot stands for root mean square of residuals. (b) Histograms of residuals. (c) Reverse cumulative distribution of absolute values of residuals. LR and polynomial SVM models have the lowest and highest fractions of isolates having high residuals, respectively. (d) Precision-recall curves. LR outperforms other models based on the residual distribution and precision-recall curve.

#### 4.3.5. MDR-hvKp pathotype

According to Table 6 and the sensitivity value for Meropenem-based models, if an isolate is resistant to Meropenem, the isolate is an MDR-hv*Kp* strain with a probability of 60-98% with a 95% C.I., Therefore, it can be concluded that we can detect MDR-hv*Kp* strains only based on Meropenem. However, precision value for the models based on Meropenem is much less than the models based on all antibiotics except Ampicillin. Thus, Meropenem-based models detect more isolates which are non-MDR-hv*Kp* as MDR-hv*Kp* strains.

**Table 6.** Performance of ML Models predicting the presence of *MDR-hvKp* in the test dataset based on sensitivity, precision, and F1-Score.

| Feature Selection method | Features | LR |  |  | NBC |  |  | LDA |  |  | SVM-Sigmoid |  |  | SVM-Polynomial |  |  | CatBoost |  |  |
| --- | --- | --- | --- | --- | --- | --- | --- | --- | --- | --- | --- | --- | --- | --- | --- | --- | --- | --- | --- |
|  |  | S (95% C.I.) | P (95% C.I.) | F1 (95% C.I.) | S (95% C.I.) | P (95% C.I.) | F1 (95% C.I.) | S (95% C.I.) | P (95% C.I.) | F1 (95% C.I.) | S (95% C.I.) | P (95% C.I.) | F1 (95% C.I.) | S (95% C.I.) | P (95% C.I.) | F1 (95% C.I.) | S (95% C.I.) | P (95% C.I.) | F1 (95% C.I.) |
| Pearson's Chi-Squared test | IMI+GN+ CIP+M<br>EM+AZ<br>M+AK+<br>CTX+FE<br>P+CAZ+<br>CRO | 0.93<br>(0.68-<br>1.00) | 0.88<br>(0.62-<br>0.98) | 0.90<br>(0.79-<br>1.00) | 0.80<br>(0.52-<br>0.96) | 0.92<br>(0.64-<br>1.00) | 0.86<br>(0.72-<br>1.00) | 0.87<br>(0.60-<br>0.98) | 0.68<br>(0.43-<br>0.87) | 0.76<br>(0.61-<br>0.92) | 0.87<br>(0.60-<br>0.98) | 0.76<br>(0.50-<br>0.93) | 0.81<br>(0.67-<br>0.96) | 1.00<br>(0.78-<br>1.00) | 0.94<br>(0.74-<br>1.00) | 0.97<br>(0.90-<br>1.00) | 0.93<br>(0.68-<br>1.00) | 0.93<br>(0.68-<br>1.00) | 0.93<br>(0.84-<br>1.00) |
| Model agnostic in all algorithms | MEM | 0.87<br>(0.60-0.98) | 0.65<br>(0.41-0.85) | 0.74<br>(0.58-0.90) | 0.87<br>(0.60-0.98) | 0.65<br>(0.41-0.85) | 0.74<br>(0.58-0.90) | 0.87<br>(0.60-0.98) | 0.65<br>(0.41-0.85) | 0.74<br>(0.58-0.90) | 0.87<br>(0.60-0.98) | 0.65<br>(0.41-0.85) | 0.74<br>(0.58-0.90) | 0.87<br>(0.60-0.98) | 0.65<br>(0.41-0.85) | 0.74<br>(0.58-0.90) | 0.87<br>(0.60-0.98) | 0.65<br>(0.41-0.85) | 0.74<br>(0.58-0.90) |

The analogy between effective antibiotics for MDR-hv*Kp* pathotypes and crucial antibiotics in hv*Kp* and MDR-c*Kp* pathotypes shows that effective antibiotics for predicting MDR-hv*Kp* strains are composed of summation of effective antibiotics in the prediction of hv*Kp* and MDR-c*Kp* except for Cefepime, which is not considered effective antibiotic for these two pathotypes. However, it is an effective antibiotic for predicting the MDR-hv*Kp* strain.

Results of feature selection based on model-agnostic approach are shown in Fig. 14. Performance of all five models based on all antibiotics, except Ampicillin, is shown in Fig. 15. As we know, plasmids play an important role in the circulation and transmission of AMR in bacteria through various plasmid-mediated conjugation mechanisms. Also, some can introduce part of genes into the bacterial chromosome or other plasmids by using high-frequency recombination (Hfr) (33). In addition to AMR plasmids in some bacteria, such as hv*Kp* isolates, there are other plasmids that encode virulence factors and are transmissible like other plasmids. In hv*Kp*s, in addition to the chromosomal virulence factors, the main virulence factors that differentiate these isolates from the classical pathotype, such as aerobactin, are encoded by large virulence plasmids (such as pLVPK and pVir) (34).

**Fig. 14.**
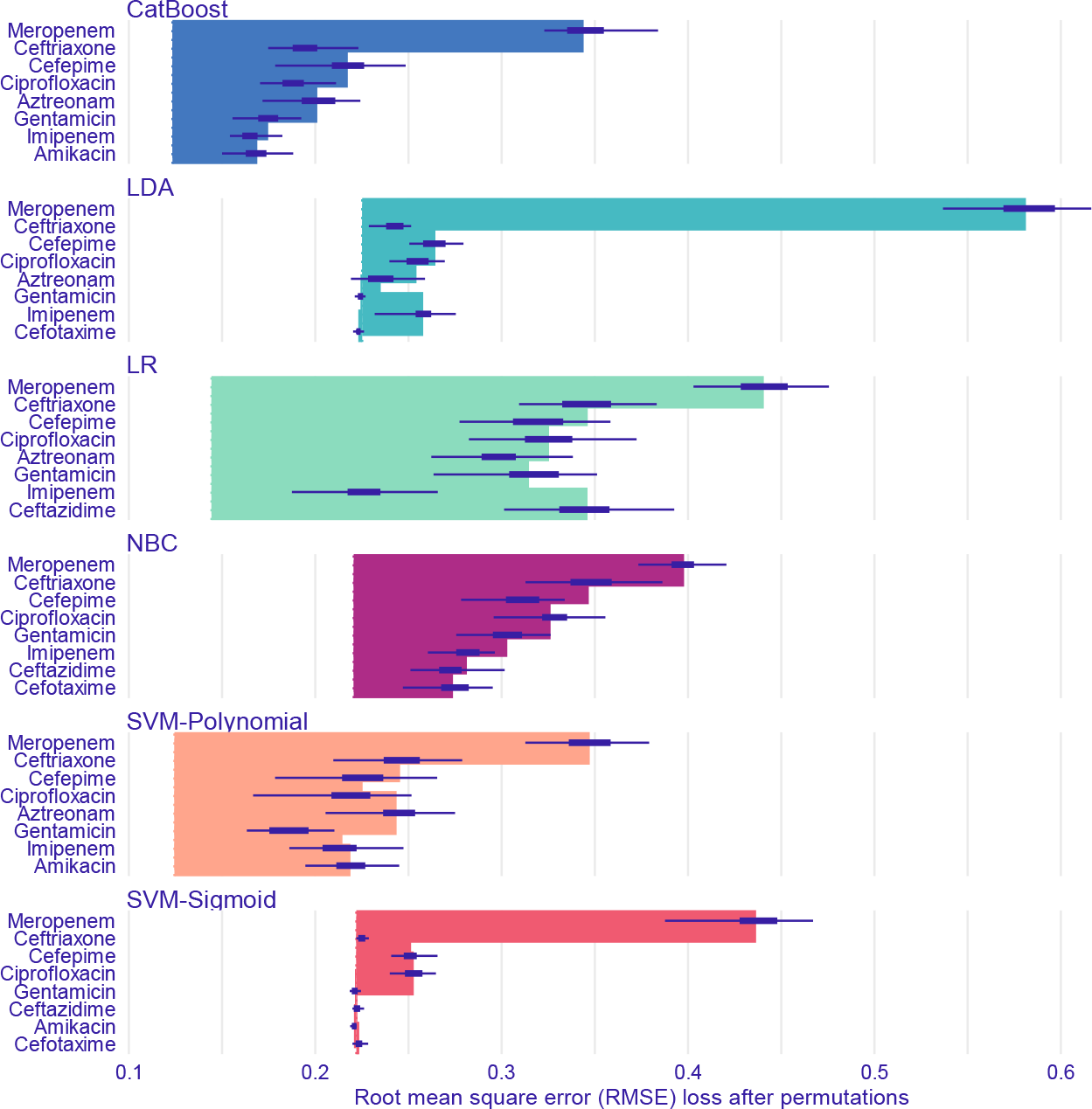
Feature importance of ML models predicting *MDR-hvKp* pathotype through model-agnostic approach over 50 permutations for top 8 antibiotics

**Fig. 15.**
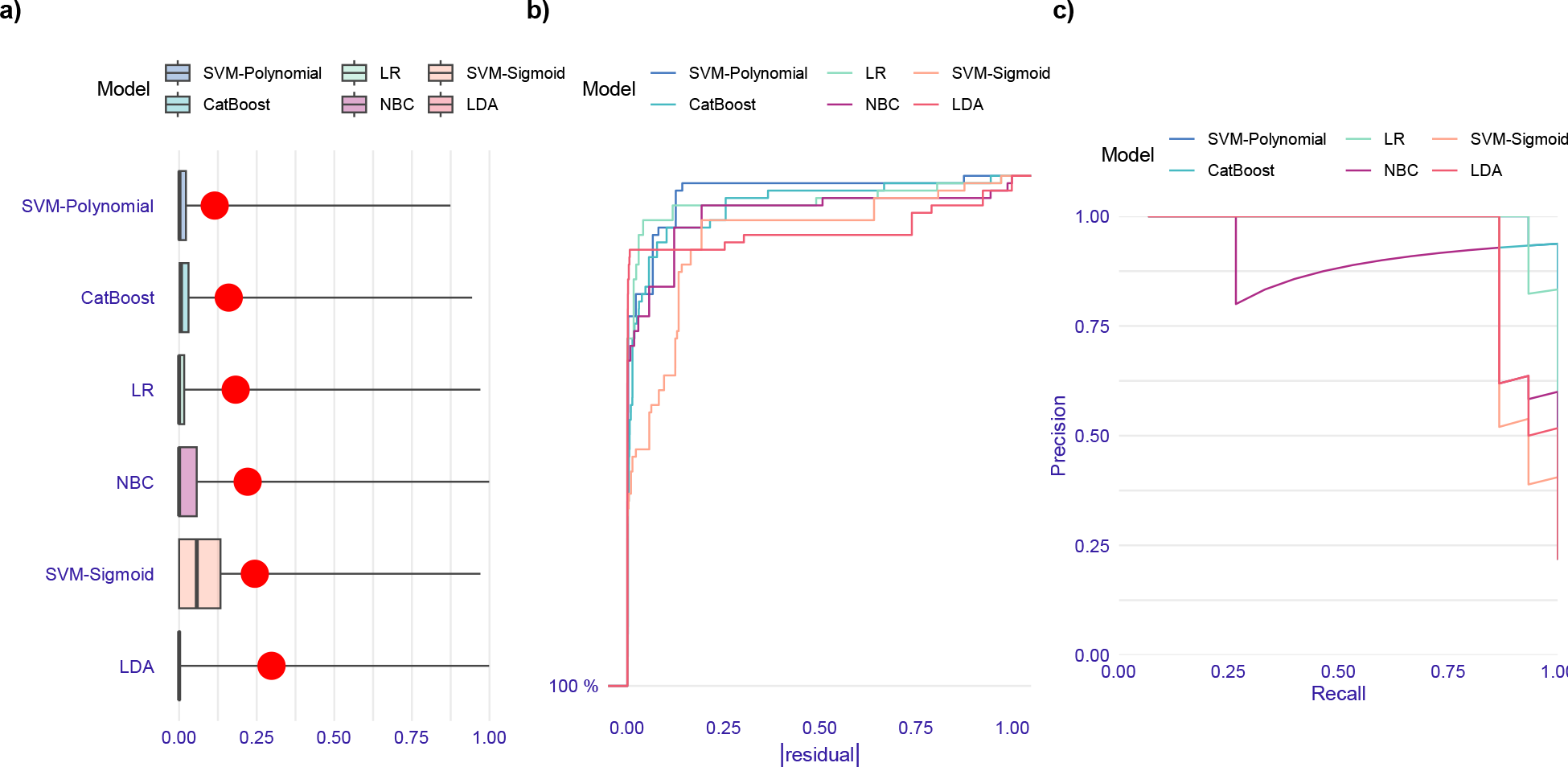
Evaluation of ML models performance in prediction of *MDR-hvKp* pathotype gene based on the test data set. (a) Boxplots of absolute values of residuals. Red dot stands for root mean square of residuals. (b) Histograms of residuals. (c) Reverse cumulative distribution of absolute values of residuals. Polynomial SVM and LDA models have the lowest and highest fractions of isolates having high residuals, respectively. (d) Precision-recall curves. SVM-polynomial model outperforms other models regarding residual distribution and precision-recall curve.

On the other hand, we know that plasmids are divided into compatible and incompatible plasmids based on their origin, and incompatible plasmids cannot coexist in a bacterial cell simultaneously and stably (35). In a valuable recent study by Yang et al. in Hong Kong, it was shown that virulence plasmids were also able to be transmitted through conjugation. The noteworthy point was that the transfer of the virulence plasmid was done by the fusion process of the virulence plasmids, with two helper plasmids including Incl1 and ColRNAI (34). Since the ColRNAI family and the Inc group play a significant role in the transfer of AMR genes, the recombination of virulence plasmids with plasmids AMR genes or their simultaneous transfer can be a logical justification for the relationship between the antibiotic resistance profile and the prediction of virulence factor genes presence in hv*Kp* isolates (36, 37).

### 4.4. ML results for beta-lactamase genetic resistance profiles prediction

#### 4.4.1. *bla*_SHV_ gene

According to table 7, by comparing performance of all ML models regarding the simplicity principle, we can conclude that for the detection of *bla*_SHV_, Cefotaxime is sufficient. Its coefficient in the LR model is positive; therefore, if an isolate is resistant to the Cefotaxime antibiotic, it carries the *bla*_SHV_ gene with a high probability of 92-100% with a 95% confidence interval. Moreover, according to the precision value, 60-85% of predictions that show isolates carrying *bla*_SHV_ are correct with a 95% confidence interval.

**Table 7.**
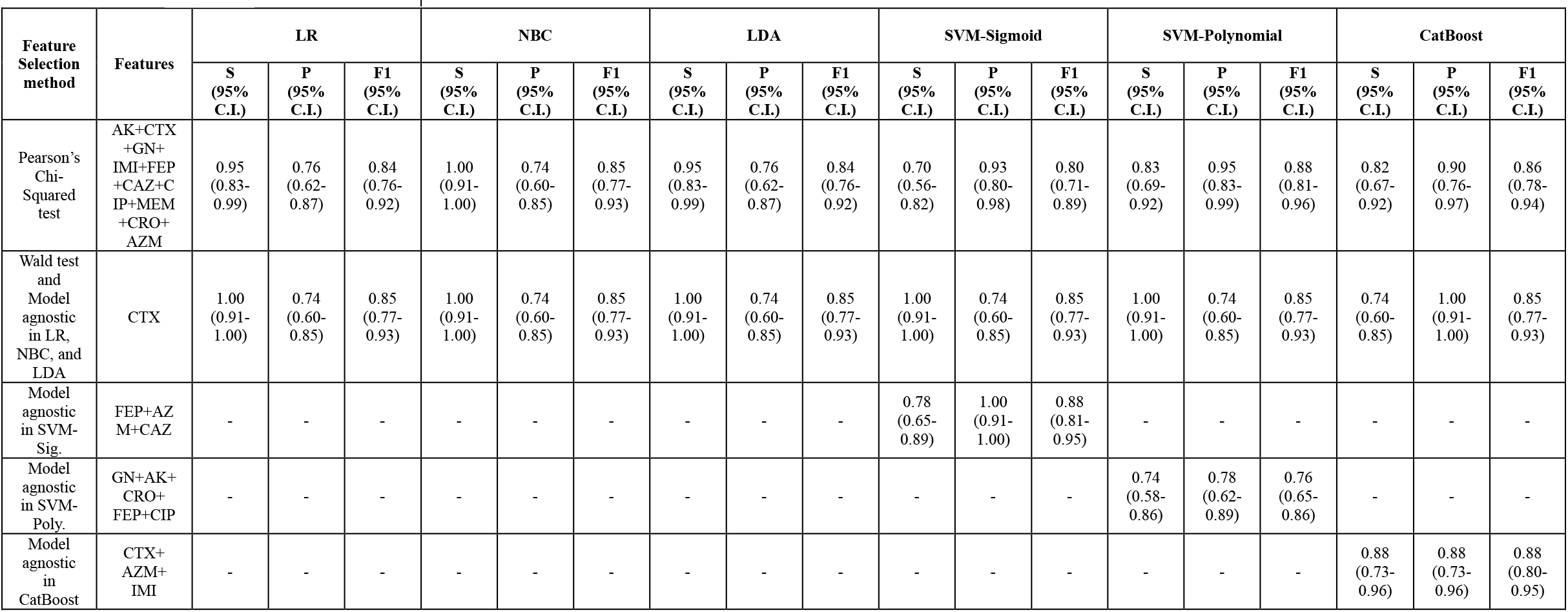
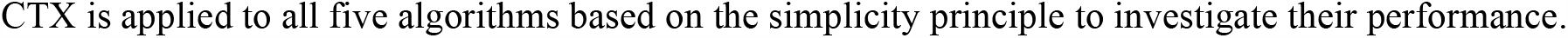
Performance of ML Models predicting the presence of *bla*_*SHV*_ in the test dataset based on sensitivity, precision, and F1-Score.

| Feature Selection method | Features | LR |  |  | NBC |  |  | LDA |  |  | SVM-Sigmoid |  |  | SVM-Polynomial |  |  | CatBoost |  |  |
| --- | --- | --- | --- | --- | --- | --- | --- | --- | --- | --- | --- | --- | --- | --- | --- | --- | --- | --- | --- |
|  |  | S (95% C.I.) | P (95% C.I.) | F1 (95% C.I.) | S (95% C.I.) | P (95% C.I.) | F1 (95% C.I.) | S (95% C.I.) | P (95% C.I.) | F1 (95% C.I.) | S (95% C.I.) | P (95% C.I.) | F1 (95% C.I.) | S (95% C.I.) | P (95% C.I.) | F1 (95% C.I.) | S (95% C.I.) | P (95% C.I.) | F1 (95% C.I.) |
| Pearson's Chi-Squared test | AK+CTX+GN+IMI+FEP+CAZ+CIP+MEM+CRO+AZM | 0.95 (0.83-0.99) | 0.76 (0.62-0.87) | 0.84 (0.76-0.92) | 1.00 (0.91-1.00) | 0.74 (0.60-0.85) | 0.85 (0.77-0.93) | 0.95 (0.83-0.99) | 0.76 (0.62-0.87) | 0.84 (0.76-0.92) | 0.70 (0.56-0.82) | 0.93 (0.80-0.98) | 0.80 (0.71-0.89) | 0.83 (0.69-0.92) | 0.95 (0.83-0.99) | 0.88 (0.81-0.96) | 0.82 (0.67-0.92) | 0.90 (0.76-0.97) | 0.86 (0.78-0.94) |
| Wald test and Model agnostic in LR, NBC, and LDA | CTX | 1.00 (0.91-1.00) | 0.74 (0.60-0.85) | 0.85 (0.77-0.93) | 1.00 (0.91-1.00) | 0.74 (0.60-0.85) | 0.85 (0.77-0.93) | 1.00 (0.91-1.00) | 0.74 (0.60-0.85) | 0.85 (0.77-0.93) | 1.00 (0.91-1.00) | 0.74 (0.60-0.85) | 0.85 (0.77-0.93) | 1.00 (0.91-1.00) | 0.74 (0.60-0.85) | 0.85 (0.77-0.93) | 0.74 (0.60-0.85) | 1.00 (0.91-1.00) | 0.85 (0.77-0.93) |
| Model agnostic in SVM-Sig. | FEP+AZM+CAZ | - | - | - | - | - | - | - | - | - | 0.78 (0.65-0.89) | 1.00 (0.91-1.00) | 0.88 (0.81-0.95) | - | - | - | - | - | - |
| Model agnostic in SVM-Poly. | GN+AK+CRO+FEP+CIP | - | - | - | - | - | - | - | - | - | - | - | - | 0.74 (0.58-0.86) | 0.78 (0.62-0.89) | 0.76 (0.65-0.86) | - | - | - |
| Model agnostic in CatBoost | CTX+AZM+IMI | - | - | - | - | - | - | - | - | - | - | - | - | - | - | - | 0.88 (0.73-0.96) | 0.88 (0.73-0.96) | 0.88 (0.80-0.95) |

CTX is applied to all five algorithms based on the simplicity principle to investigate their performance.

Results of feature (antibiotics) importance analysis through the model-agnostic approach are shown in Fig. 16. Additionally, performance of all five algorithms based on all antibiotics, except Ampicillin, in terms of residual distribution and precision-recall curve are represented in Fig. 17

**Fig. 16.**
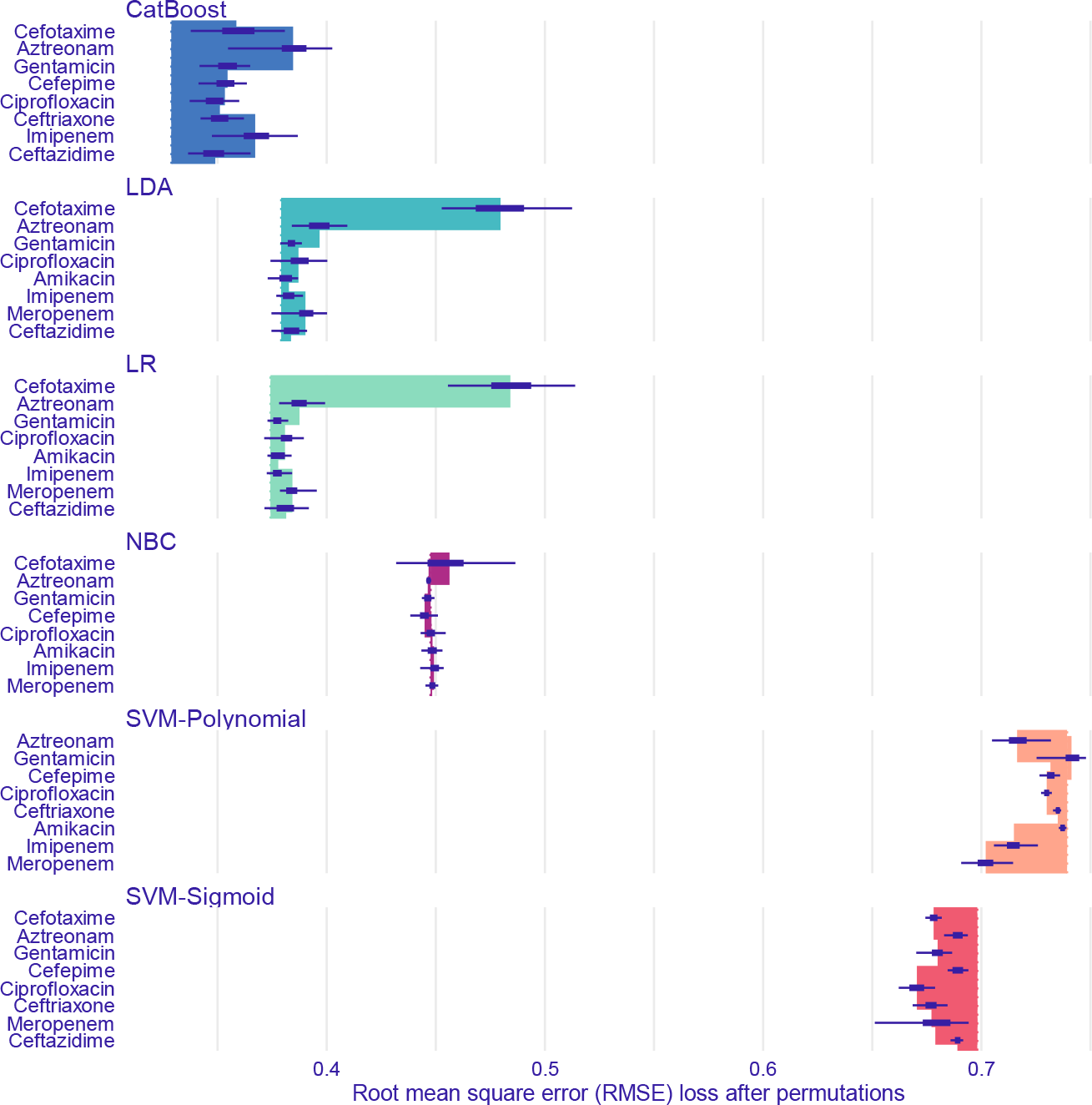
Feature importance of ML models predicting *bla*_SHV_ gene through model-agnostic approach over 50 permutations for top 8 antibiotics

**Fig. 17.**
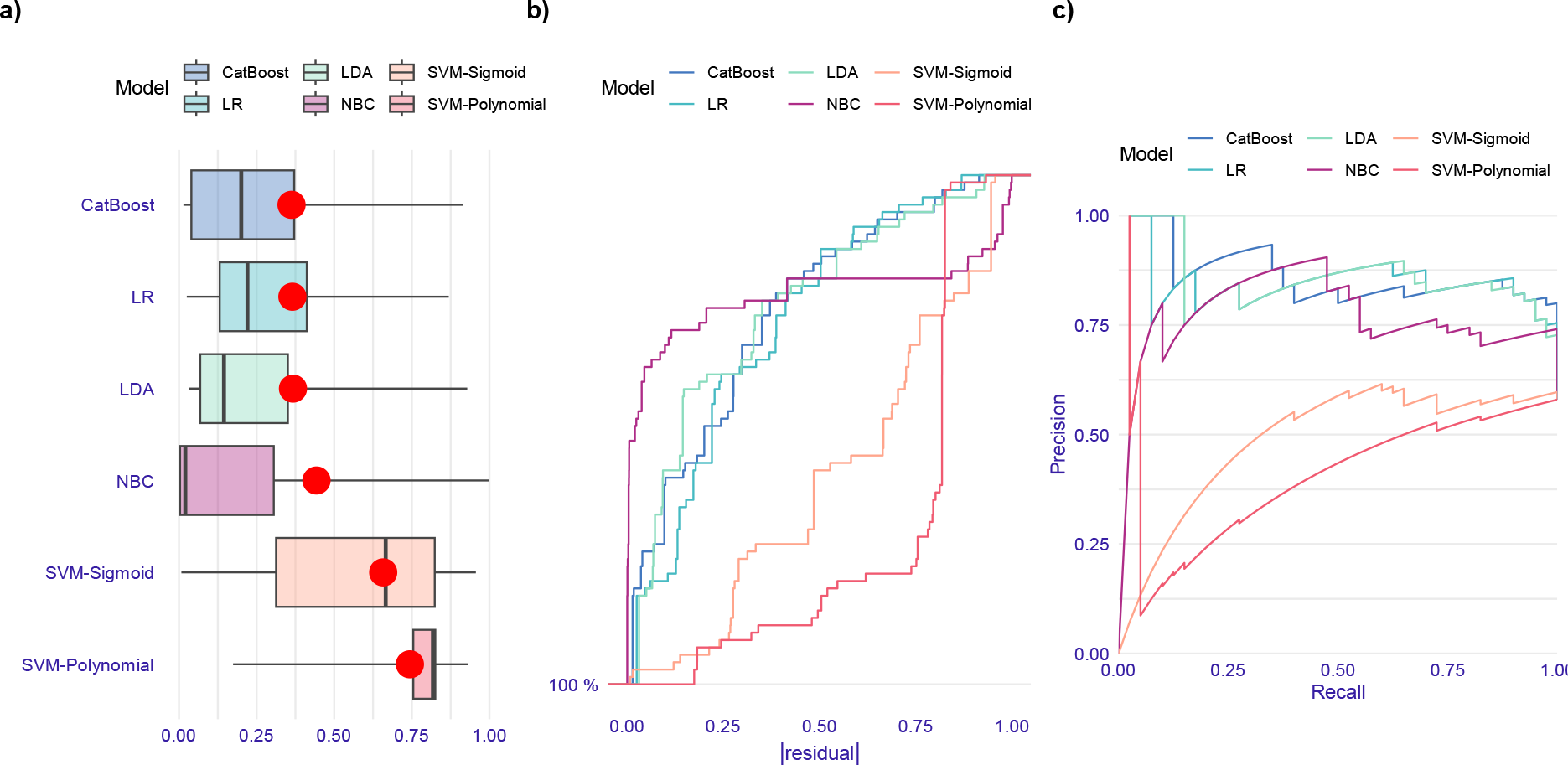
Evaluation of ML models performance in prediction of *bla*_SHV_ gene based on the test data set. (a) Boxplots of absolute values of residuals. Red dot stands for root mean square of residuals. (b) Reverse cumulative distribution of absolute values of residuals. Polynomial SVM model has the highest fraction and CatBoost, LR, and LDA models have the lowest fractions of the isolates having high residuals. Sigmoid SVM model has more uniform distribution of residuals than other models. (c) Precision-recall curves. CatBoost, LDA, and LR models have similar performance and outperform other ML models.

Cefotaxime, a third-generation cephalosporin was first synthesized in 1976. Despite the broad-spectrum bactericidal activity of this antibiotic against gram-negative and gram-positive bacterial infections, *K. pneumoniae* isolates carrying CTX-type beta-lactamase were identified in France in the late-1980s (40, 41). The studies that investigated the relationship between cefotaxime antibiotic resistance and the presence of different types of the *bla*_SHV_ gene, have revealed that *bla*_SHV_ carrier isolates often show resistance to the cefotaxime antibiotic. Consequently, the ML analysis findings in this paper are in line with the other studies in the field of microbiology (42, 43).

#### 4.4.2. *bla*_TEM_

According to Table 8, ML models based on Cefotaxime have the highest sensitivity. Its coefficient in the logistic function is positive; if an isolate is resistant to the Cefotaxime, the isolate carries *bla*_TEM_ with a probability of 89-100% with a 95% confidence interval. Therefore, we can only consider Cefotaxime to detect the presence of *bla*_TEM_. Meanwhile, only 47-74% of these predictions are correct with a 95% C.I, which means we can detect the presence of the gene confidently (High sensitivity), but 26-53% of the isolates not carrying the gene are misclassified as isolates carrying *bla*_TEM_.

**Table 8.**
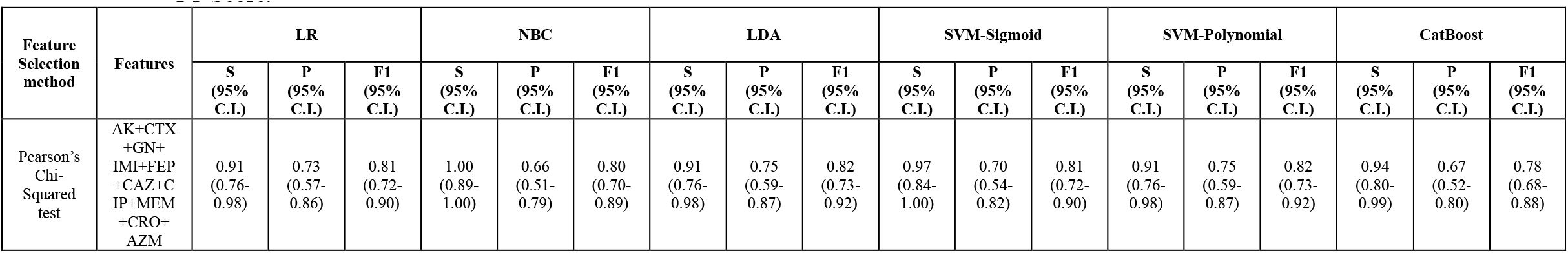
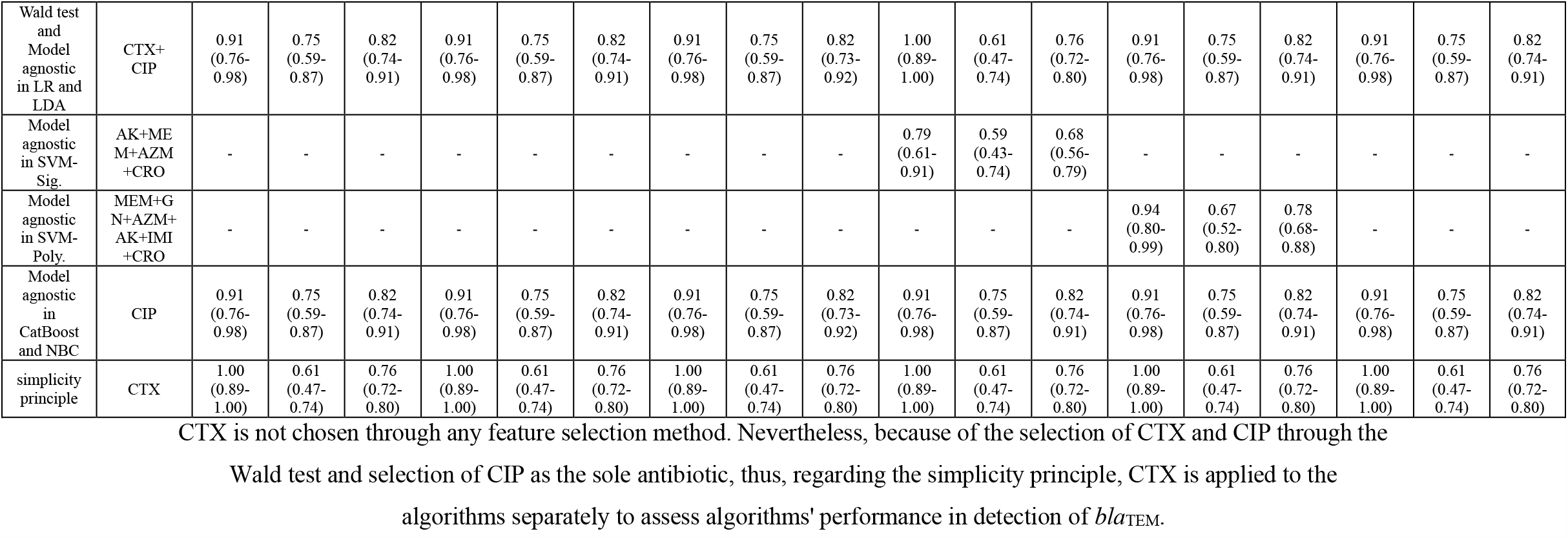
Performance of ML Models predicting the presence of *bla*_*TEM*_ in the test dataset based on sensitivity, precision, and F1-Score.

Results of feature (antibiotic) importance analysis by the model-agnostic approach for all the algorithms are represented in Fig. 18 and performance of all models considering residual distribution and precision-recall curve are shown in Fig. 19 Based on all antibiotics except Ampicillin and

**Fig. 18.**
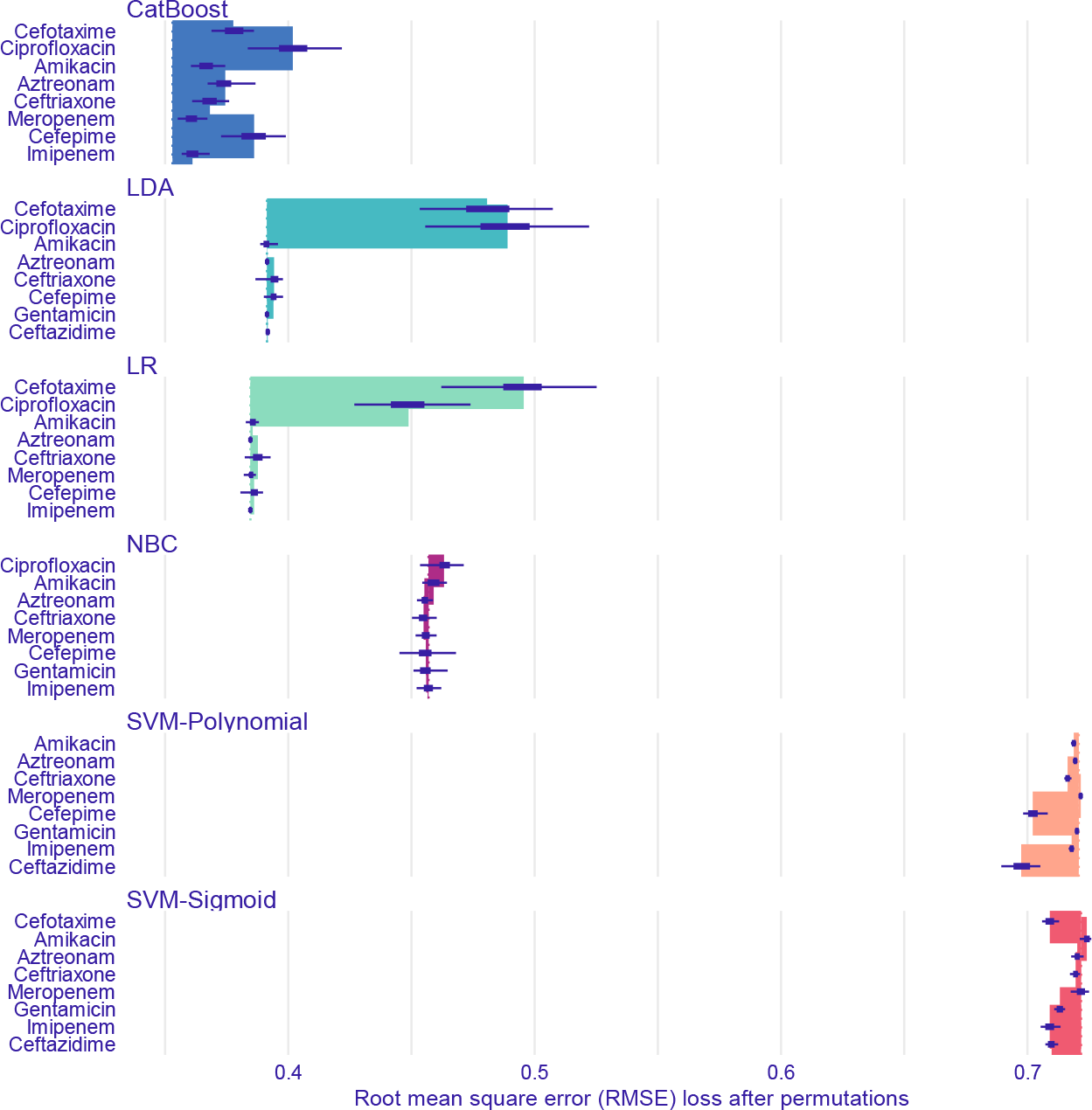
Feature importance of ML models predicting *bla*_TEM_ gene through model-agnostic approach over 50 permutations for top 8 antibiotics

**Fig. 19.**
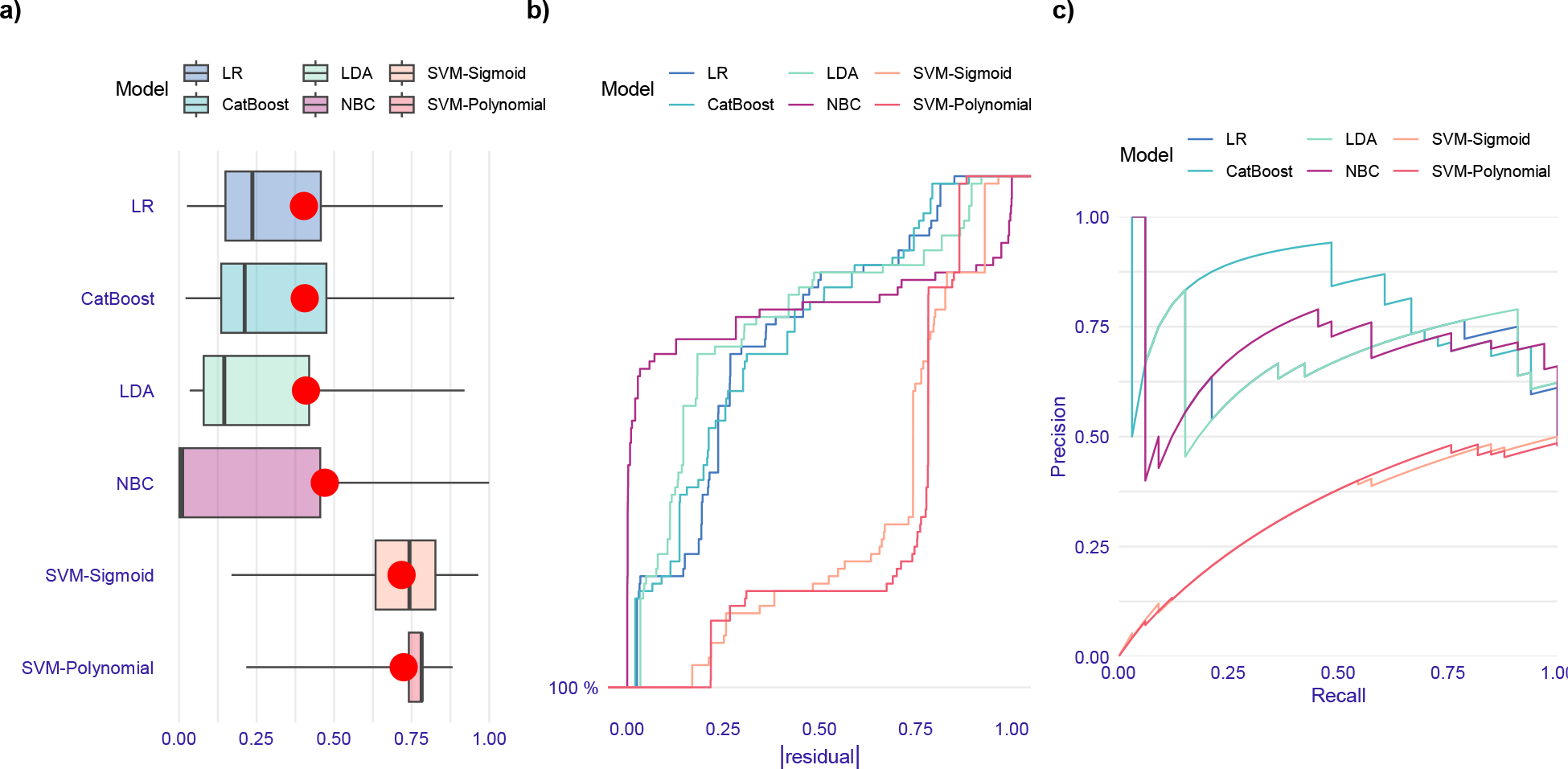
Evaluation of ML models performance in prediction of *bla*_TEM_ gene based on the test data set. (a) Boxplots of absolute values of residuals. Red dot stands for root mean square of residuals. (b) Reverse cumulative distribution of absolute values of residuals. LR, CatBoost, and LDA models have the lowest fractions of residuals and polynomial SVM model has the highest fraction. Distribution of residuals in sigmoid SVM model is more uniform than other models (d) Precision-recall curves. Performance of CatBoost, LR, and LDA are close to each other and they outperform the others based on the precision-recall curve and residuals distributions.

#### 4.4.3. *bla*_CTX-M15_ gene

According to Table 9, despite the higher sensitivity of Cefotaxime-based models than models based on Cefepime, it is a negligible difference, and Cefotaxime-based ML models show much less precision and consequently, less F1-Score than Cefepime-based models. Therefore, Cefepime-based models outperform Cefotaxime-based ones. Also, considering the simplicity principle, the model based on the Cefepime antibiotic is the optimal option. Since the positive coefficient of FEP in its logistic function, if the isolate shows resistance to the Cefepime antibiotic, it carries a *bla*_CTX-M15_ gene with a probability of 80-98% with a 95% confidence interval. Also, concerning the precision value, 72-95% of predictions showing the presence of the gene are correct. Moreover, if an isolate is resistant to Cefotaxime, it carries a *bla*_CTX-M15_ gene with a probability of 87-100% with a 95% confidence interval. This means the probability of detecting isolates carrying *bla*_CTX-M15_ based on Cefotaxime is higher than Cefepime.

**Table 9.**
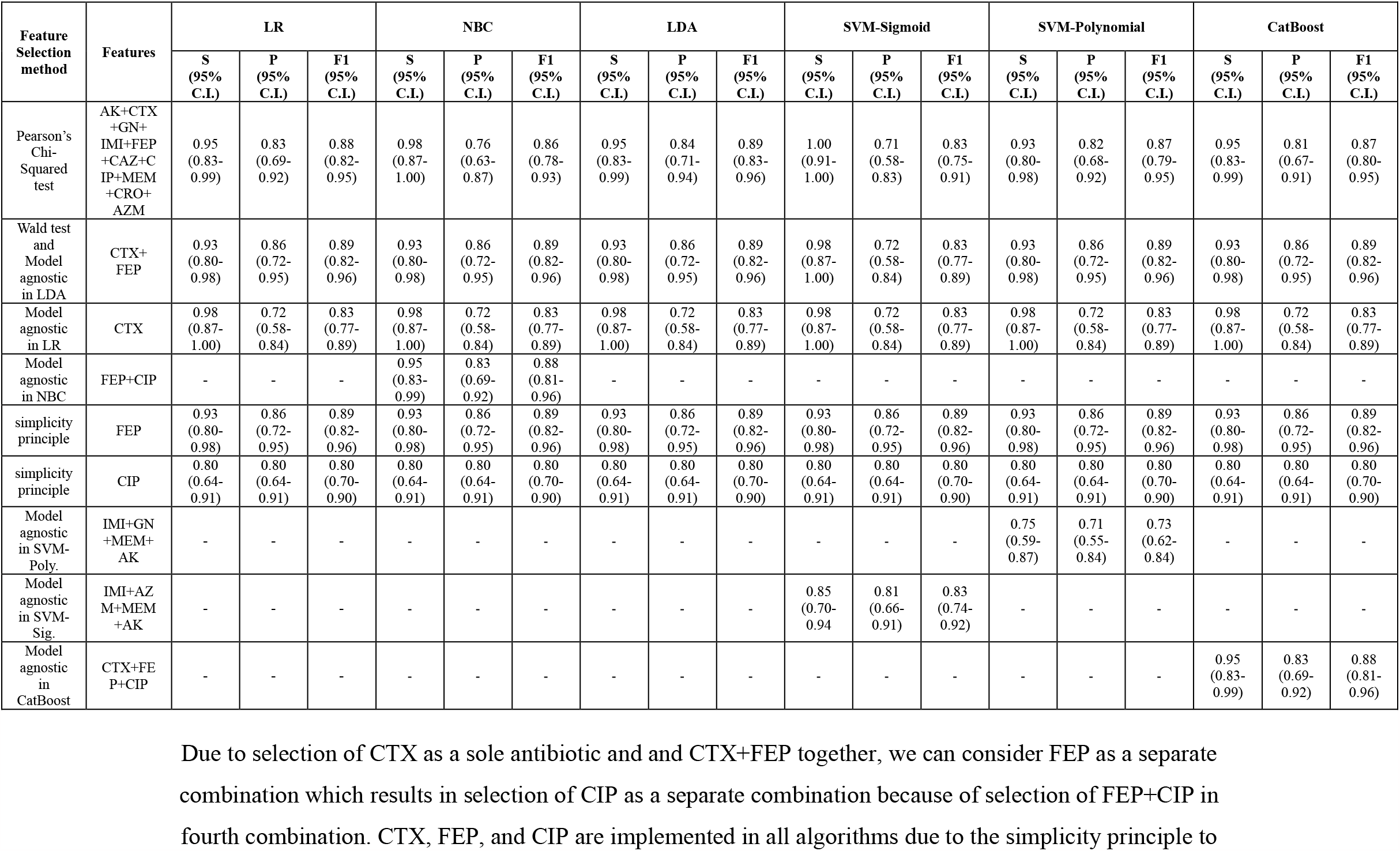
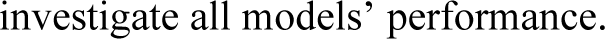
Performance of ML Models predicting the presence of *bla*_*CTX-M15*_ in test dataset based on sensitivity, precision, and F1-Score.

| Feature Selection method | Features | LR |  |  | NBC |  |  | LDA |  |  | SVM-Sigmoid |  |  | SVM-Polynomial |  |  | CatBoost |  |  |
| --- | --- | --- | --- | --- | --- | --- | --- | --- | --- | --- | --- | --- | --- | --- | --- | --- | --- | --- | --- |
|  |  | S (95% C.I.) | P (95% C.I.) | F1 (95% C.I.) | S (95% C.I.) | P (95% C.I.) | F1 (95% C.I.) | S (95% C.I.) | P (95% C.I.) | F1 (95% C.I.) | S (95% C.I.) | P (95% C.I.) | F1 (95% C.I.) | S (95% C.I.) | P (95% C.I.) | F1 (95% C.I.) | S (95% C.I.) | P (95% C.I.) | F1 (95% C.I.) |
| Pearson's Chi-Squared test | AK+CTX+GN+IMI+FEP+CAZ+CIP+MEM+CRO+AZM | 0.95 (0.83-0.99) | 0.83 (0.69-0.92) | 0.88 (0.82-0.95) | 0.98 (0.87-1.00) | 0.76 (0.63-0.87) | 0.86 (0.78-0.93) | 0.95 (0.83-0.99) | 0.84 (0.71-0.94) | 0.89 (0.83-0.96) | 1.00 (0.91-1.00) | 0.71 (0.58-0.83) | 0.83 (0.75-0.91) | 0.93 (0.80-0.98) | 0.82 (0.68-0.92) | 0.87 (0.79-0.95) | 0.95 (0.83-0.99) | 0.81 (0.67-0.91) | 0.87 (0.80-0.95) |
| Wald test and Model agnostic in LDA | CTX+FEP | 0.93 (0.80-0.98) | 0.86 (0.72-0.95) | 0.89 (0.82-0.96) | 0.93 (0.80-0.98) | 0.86 (0.72-0.95) | 0.89 (0.82-0.96) | 0.93 (0.80-0.98) | 0.86 (0.72-0.95) | 0.89 (0.82-0.96) | 0.98 (0.87-1.00) | 0.72 (0.58-0.84) | 0.83 (0.77-0.89) | 0.93 (0.80-0.98) | 0.86 (0.72-0.95) | 0.89 (0.82-0.96) | 0.93 (0.80-0.98) | 0.86 (0.72-0.95) | 0.89 (0.82-0.96) |
| Model agnostic in LR | CTX | 0.98 (0.87-1.00) | 0.72 (0.58-0.84) | 0.83 (0.77-0.89) | 0.98 (0.87-1.00) | 0.72 (0.58-0.84) | 0.83 (0.77-0.89) | 0.98 (0.87-1.00) | 0.72 (0.58-0.84) | 0.83 (0.77-0.89) | 0.98 (0.87-1.00) | 0.72 (0.58-0.84) | 0.83 (0.77-0.89) | 0.98 (0.87-1.00) | 0.72 (0.58-0.84) | 0.83 (0.77-0.89) | 0.98 (0.87-1.00) | 0.72 (0.58-0.84) | 0.83 (0.77-0.89) |
| Model agnostic in NBC | FEP+CIP | - | - | - | 0.95 (0.83-0.99) | 0.83 (0.69-0.92) | 0.88 (0.81-0.96) | - | - | - | - | - | - | - | - | - | - | - | - |
| simplicity principle | FEP | 0.93 (0.80-0.98) | 0.86 (0.72-0.95) | 0.89 (0.82-0.96) | 0.93 (0.80-0.98) | 0.86 (0.72-0.95) | 0.89 (0.82-0.96) | 0.93 (0.80-0.98) | 0.86 (0.72-0.95) | 0.89 (0.82-0.96) | 0.93 (0.80-0.98) | 0.86 (0.72-0.95) | 0.89 (0.82-0.96) | 0.93 (0.80-0.98) | 0.86 (0.72-0.95) | 0.89 (0.82-0.96) | 0.93 (0.80-0.98) | 0.86 (0.72-0.95) | 0.89 (0.82-0.96) |
| simplicity principle | CIP | 0.80 (0.64-0.91) | 0.80 (0.64-0.91) | 0.80 (0.70-0.90) | 0.80 (0.64-0.91) | 0.80 (0.64-0.91) | 0.80 (0.70-0.90) | 0.80 (0.64-0.91) | 0.80 (0.64-0.91) | 0.80 (0.70-0.90) | 0.80 (0.64-0.91) | 0.80 (0.64-0.91) | 0.80 (0.70-0.90) | 0.80 (0.64-0.91) | 0.80 (0.64-0.91) | 0.80 (0.70-0.90) | 0.80 (0.64-0.91) | 0.80 (0.64-0.91) | 0.80 (0.70-0.90) |
| Model agnostic in SVM-Poly. | IMI+GN+MEM+AK | - | - | - | - | - | - | - | - | - | - | - | - | 0.75 (0.59-0.87) | 0.71 (0.55-0.84) | 0.73 (0.62-0.84) | - | - | - |
| Model agnostic in SVM-Sig. | IMI+AZM+MEM+AK | - | - | - | - | - | - | - | - | - | 0.85 (0.70-0.94) | 0.81 (0.66-0.91) | 0.83 (0.74-0.92) | - | - | - | - | - | - |
| Model agnostic in CatBoost | CTX+FEP+CIP | - | - | - | - | - | - | - | - | - | - | - | - | - | - | - | 0.95 (0.83-0.99) | 0.83 (0.69-0.92) | 0.88 (0.81-0.96) |

Due to selection of CTX as a sole antibiotic and and CTX+FEP together, we can consider FEP as a separate combination which results in selection of CIP as a separate combination because of selection of FEP+CIP in fourth combination. CTX, FEP, and CIP are implemented in all algorithms due to the simplicity principle to
investigate all models' performance.

Comparing crucial antibiotics in detection of presence of *bla*_SHV_, *bla*_TEM_, and *bla*_CTX-M15_ shows that Cefotaxime plays an important role which is in compliance with the experiments showing simultaneous presence of these genes (38, 39).

Results of model-agnostic approach for antibiotics importance analysis in all algorithms are represented in Fig. 20. Moreover, regarding residual distribution and precision-recall curve, performance of all five models based on all antibiotics, except Ampicillin, are evaluated and shown in Fig. 21.

**Fig. 20.**
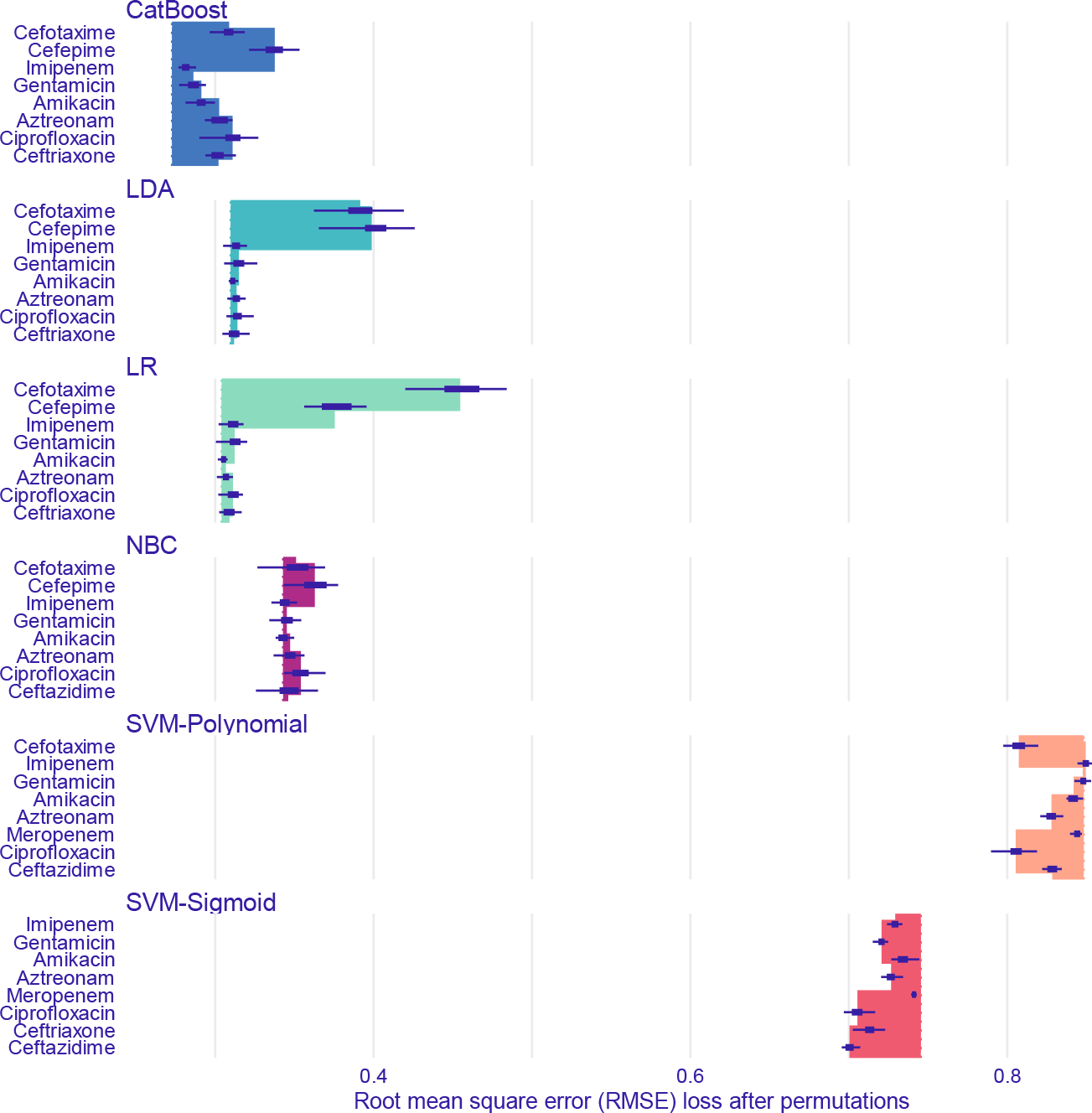
Feature importance of ML models predicting *bla*_CTX-M15_ gene through model-agnostic approach over 50 permutations for top 8 antibiotics

**Fig. 21.**
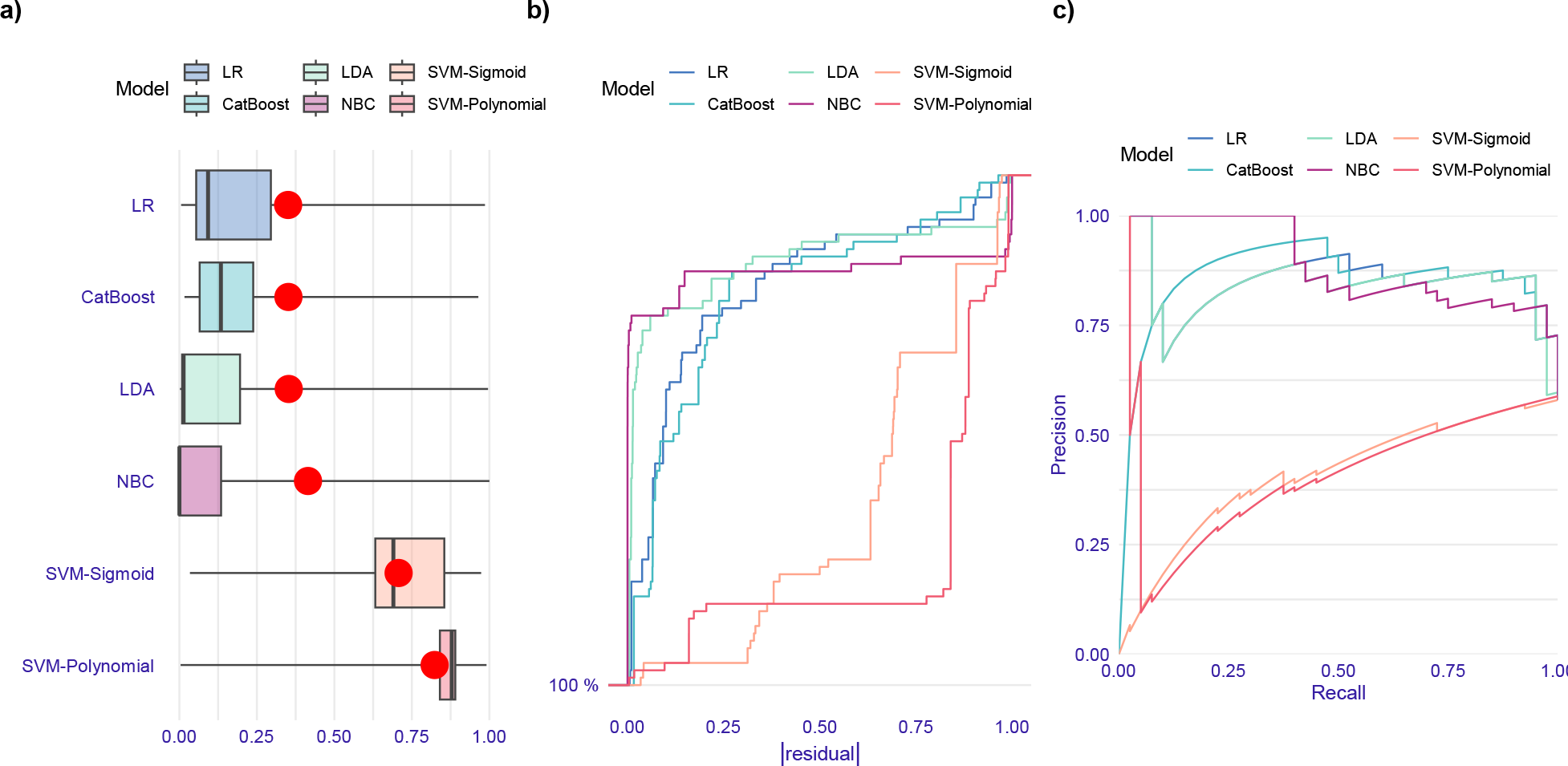
Evaluation of ML models performance in prediction of *bla*_CTX-M15_ gene based on the test data set. (a) Boxplots of absolute values of residuals. Red dot stands for root mean square of residuals. (b) Reverse cumulative distribution of absolute values of residuals. LR, LDA, and CatBoost models approximately have the same fraction of residuals and outperform other models. On the contrary, SVM polynomial has the worst performance based on the fraction of residuals. (d) Precision-recall curves. LR, LDA, and CatBoost models approximately have the same performance and they outperform other models.

#### 4.4.4. *bla*_NDM-1_ gene

According to Table 10 and considering the positive coefficient of Imipenem in Logistic function, the results of all Imipenem-based models indicated that if an isolate shows resistance to the Imipenem antibiotic; the isolate carries *bla*_NDM**-1**_ gene with a probability of 81-100% with a 95% C.I. (sensitivity), therefore, Imipenem is sufficient for detection of isolates carrying *bla*_NDM**-1**_. Although ML models based on Imipenem are much more straightforward, the NBC model based on Imipenem, Cefepime, and Ciprofloxacin has much better performance regarding precision and F1-score values. Regarding the precision value for the NBC model; 53-90% of predictions indicating the presence of *bla*_NDM-1_ based on Imipenem, Cefepime, and Ciprofloxacin, are correct.

**Table 10.**
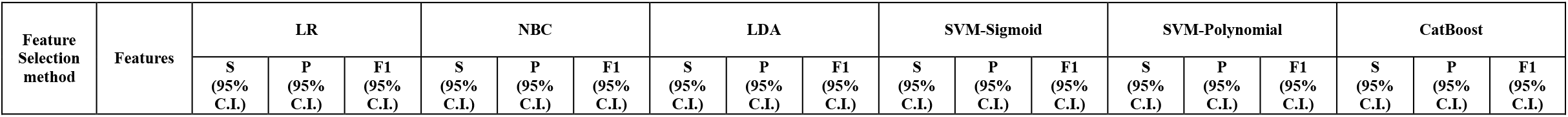
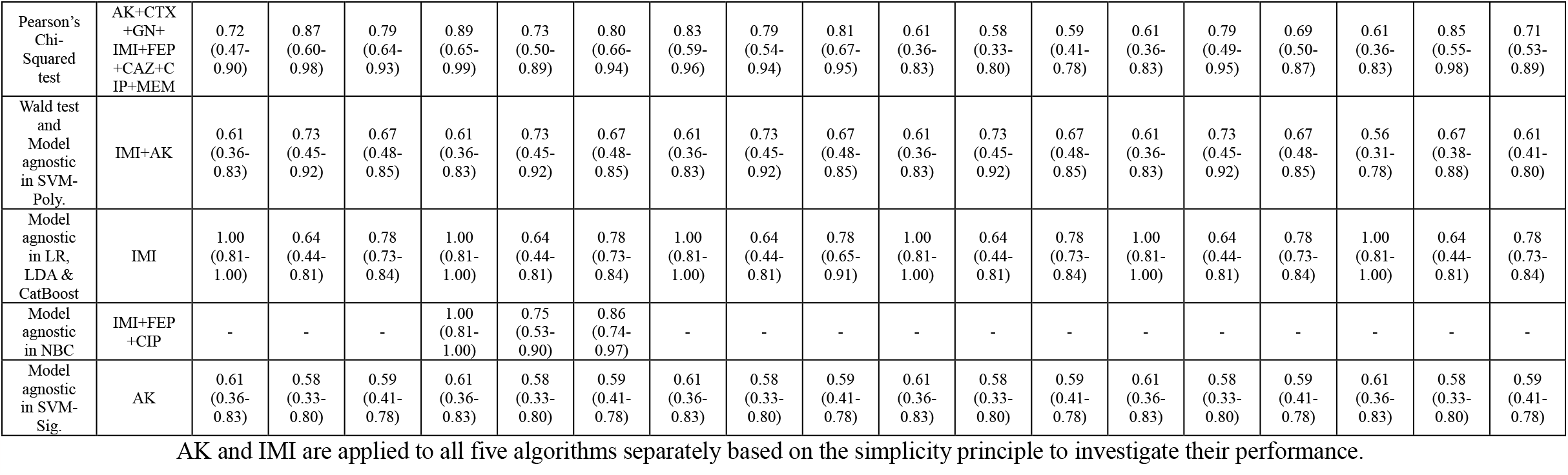
Performance of ML Models predicting the presence of *bla*_*NDM-1*_ in the test dataset based on sensitivity, precision, and F1-Score.

| Feature Selection method | Features | LR |  |  | NBC |  |  | LDA |  |  | SVM-Sigmoid |  |  | SVM-Polynomial |  |  | CatBoost |  |  |
| --- | --- | --- | --- | --- | --- | --- | --- | --- | --- | --- | --- | --- | --- | --- | --- | --- | --- | --- | --- |
|  |  | S (95% C.I.) | P (95% C.I.) | F1 (95% C.I.) | S (95% C.I.) | P (95% C.I.) | F1 (95% C.I.) | S (95% C.I.) | P (95% C.I.) | F1 (95% C.I.) | S (95% C.I.) | P (95% C.I.) | F1 (95% C.I.) | S (95% C.I.) | P (95% C.I.) | F1 (95% C.I.) | S (95% C.I.) | P (95% C.I.) | F1 (95% C.I.) |
| Pearson's Chi-Squared test | AK+CTX+GN+IMI+FEP+CAZ+CIP+MEM | 0.72<br>(0.47-0.90) | 0.87<br>(0.60-0.98) | 0.79<br>(0.64-0.93) | 0.89<br>(0.65-0.99) | 0.73<br>(0.50-0.89) | 0.80<br>(0.66-0.94) | 0.83<br>(0.59-0.96) | 0.79<br>(0.54-0.94) | 0.81<br>(0.67-0.95) | 0.61<br>(0.36-0.83) | 0.58<br>(0.33-0.80) | 0.59<br>(0.41-0.78) | 0.61<br>(0.36-0.83) | 0.79<br>(0.49-0.95) | 0.69<br>(0.50-0.87) | 0.61<br>(0.36-0.83) | 0.85<br>(0.55-0.98) | 0.71<br>(0.53-0.89) |
| Wald test and Model agnostic in SVM-Poly. | IMI+AK | 0.61<br>(0.36-0.83) | 0.73<br>(0.45-0.92) | 0.67<br>(0.48-0.85) | 0.61<br>(0.36-0.83) | 0.73<br>(0.45-0.92) | 0.67<br>(0.48-0.85) | 0.61<br>(0.36-0.83) | 0.73<br>(0.45-0.92) | 0.67<br>(0.48-0.85) | 0.61<br>(0.36-0.83) | 0.73<br>(0.45-0.92) | 0.67<br>(0.48-0.85) | 0.61<br>(0.36-0.83) | 0.73<br>(0.45-0.92) | 0.67<br>(0.48-0.85) | 0.56<br>(0.31-0.78) | 0.67<br>(0.38-0.88) | 0.61<br>(0.41-0.80) |
| Model agnostic in LR, LDA & CatBoost | IMI | 1.00<br>(0.81-1.00) | 0.64<br>(0.44-0.81) | 0.78<br>(0.73-0.84) | 1.00<br>(0.81-1.00) | 0.64<br>(0.44-0.81) | 0.78<br>(0.73-0.84) | 1.00<br>(0.81-1.00) | 0.64<br>(0.44-0.81) | 0.78<br>(0.65-0.91) | 1.00<br>(0.81-1.00) | 0.64<br>(0.44-0.81) | 0.78<br>(0.73-0.84) | 1.00<br>(0.81-1.00) | 0.64<br>(0.44-0.81) | 0.78<br>(0.73-0.84) | 1.00<br>(0.81-1.00) | 0.64<br>(0.44-0.81) | 0.78<br>(0.73-0.84) |
| Model agnostic in NBC | IMI+FEP+CIP | - | - | - | 1.00<br>(0.81-1.00) | 0.75<br>(0.53-0.90) | 0.86<br>(0.74-0.97) | - | - | - | - | - | - | - | - | - | - | - | - |
| Model agnostic in SVM-Sig. | AK | 0.61<br>(0.36-0.83) | 0.58<br>(0.33-0.80) | 0.59<br>(0.41-0.78) | 0.61<br>(0.36-0.83) | 0.58<br>(0.33-0.80) | 0.59<br>(0.41-0.78) | 0.61<br>(0.36-0.83) | 0.58<br>(0.33-0.80) | 0.59<br>(0.41-0.78) | 0.61<br>(0.36-0.83) | 0.58<br>(0.33-0.80) | 0.59<br>(0.41-0.78) | 0.61<br>(0.36-0.83) | 0.58<br>(0.33-0.80) | 0.59<br>(0.41-0.78) | 0.61<br>(0.36-0.83) | 0.58<br>(0.33-0.80) | 0.59<br>(0.41-0.78) |

AK and IMI are applied to all five algorithms separately based on the simplicity principle to investigate their performance.

Beta-lactamase type *bla*_NDM**-1**_ is a broad-spectrum carbapenem that can hydrolyze all beta-lactamase agents except aztreonam, thus, the bacterial isolates carrying *bla*_NDM**-1**_ are usually reported to be resistant to all beta-lactamase (40). Since the *bla*_NDM**-1**_ gene is carried by a plasmid and is easily transferred to other bacteria and causes the emergence of MDR-bacterial strains, diagnosis and treatment of infection caused by *bla*_NDM**-1**_-producing strains are very important in public health (41). In a study reported from Kuwait by Jamal *et al*., 21 isolates were *bla*_NDM**-1**_-producing and 14 (66.6%) isolates were identified as *bla*_NDM**-1**_ positive *K*.*pneumoniae* out of 61 isolates of carbapenem-resistant *Enterobacteriaceae* (CRE) (41). In another study, all 57 CRE isolates were known as *bla*_NDM_ producers and 22 of them were *K*.*pneumoniae* isolates (46). In 44 isolates, the *bla*_NDM**-1**_ gene (20 *K*.*pneumoniae*) was identified and 13 isolates carried other *bla*_NDM_ variants (2 *K*.*pneumoniae*) (42). Therefore, through the summary of the ML-analysis and the results of the microbiology investigations, we reached a common conclusion that the prevalence of the *bla*_NDM**-1**_ gene in carbapenem-resistant isolates was reported as high as expected.

Feature importance results based on applying model-agnostic approach to all the algorithms are shown in Fig. 22. Performance of all the algorithms based on the selected antibiotics through Pearson’s chi-squared test considering residual distribution and precision-recall curve are shown in Fig. 23.

**Fig. 22.**
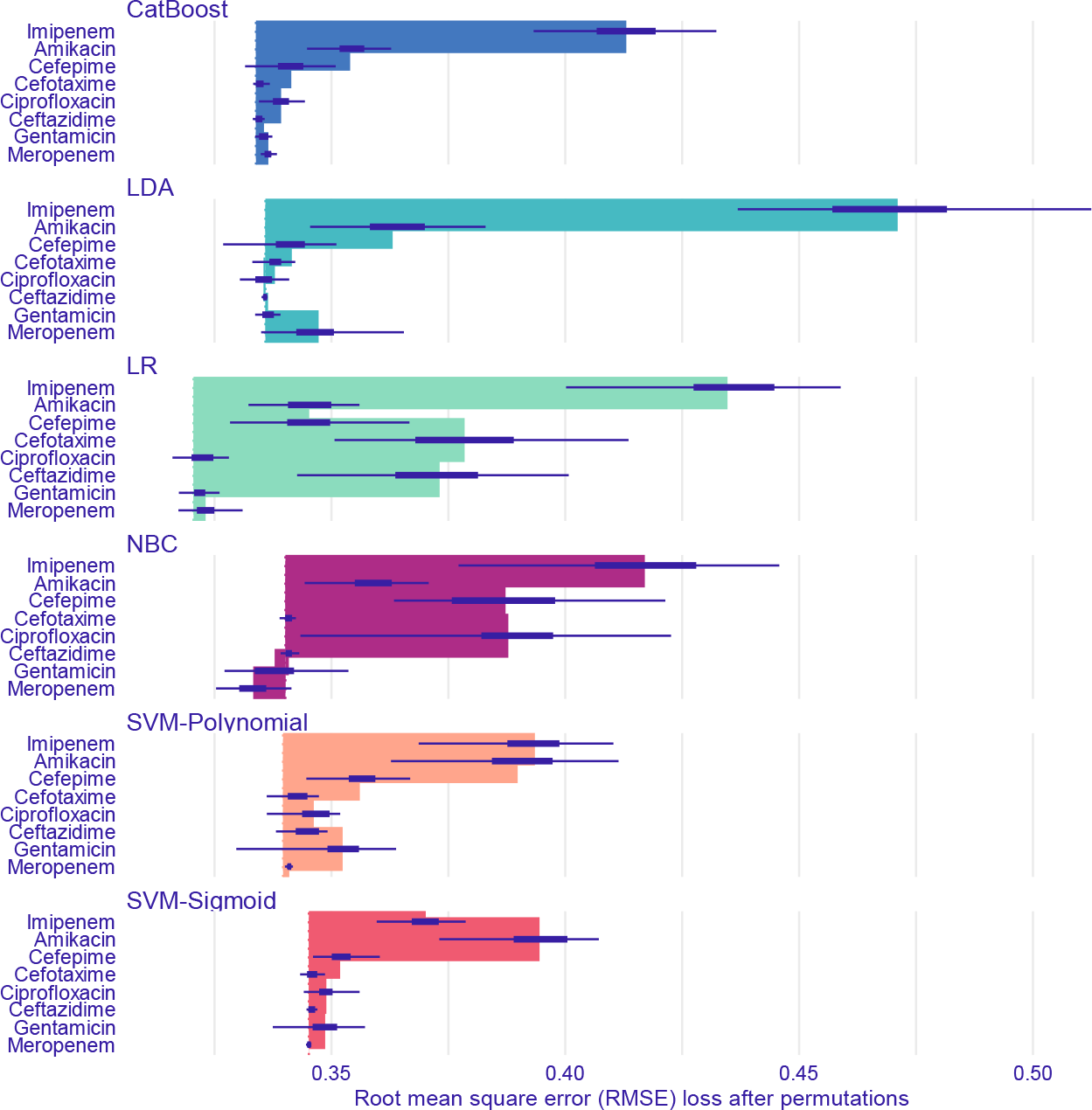
Feature importance of ML models predicting *bla*_NDM-1_ gene through model-agnostic approach over 50 permutations for top 8 antibiotics

**Fig. 23.**
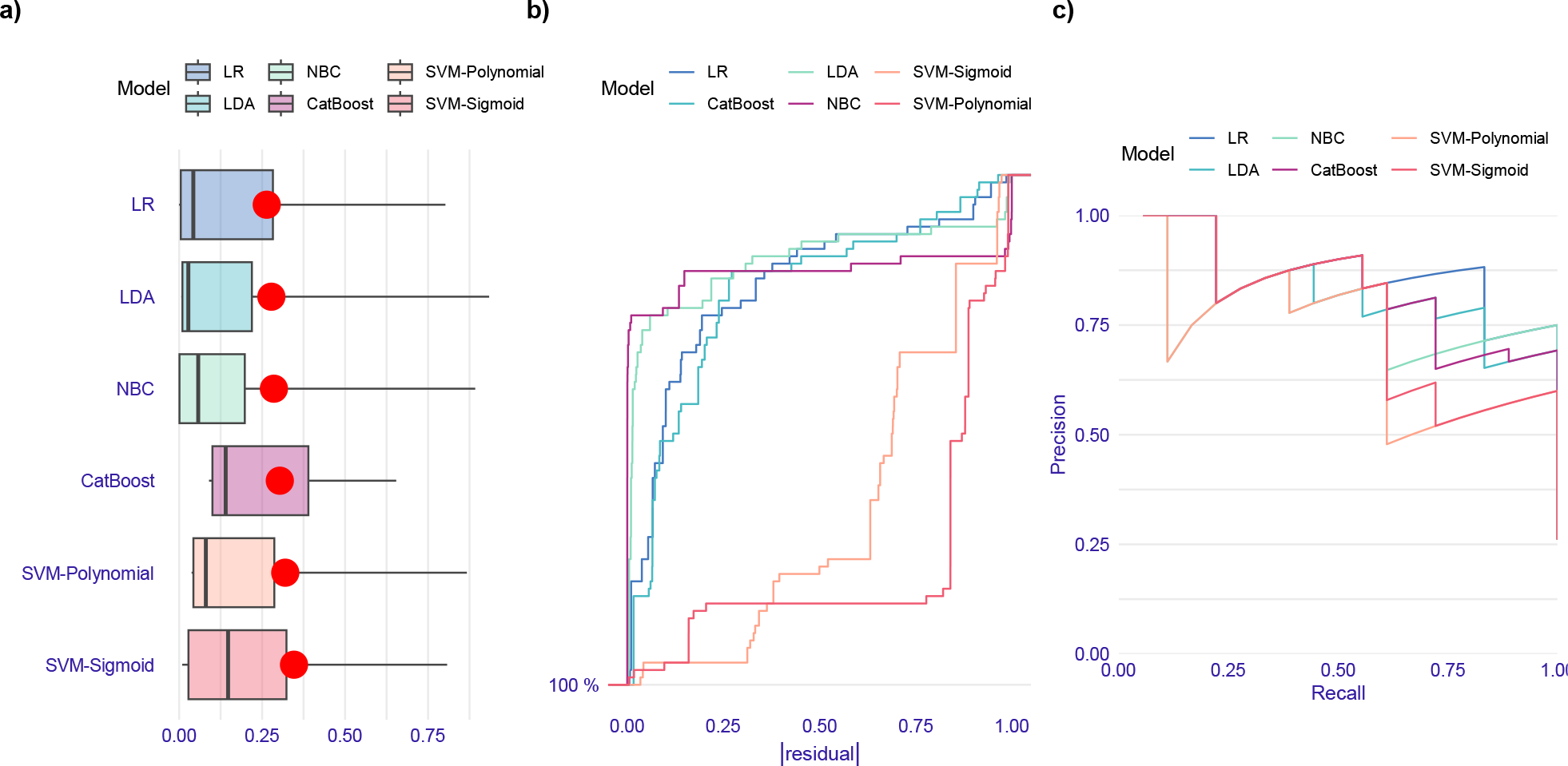
Evaluation of ML models performance in prediction of *bla*_NDM-1_ gene based on the test data set. (a) Boxplots of absolute values of residuals. Red dot stands for root mean square of residuals. (b) Reverse cumulative distribution of absolute values of residuals. LR, LDA, and NBC models have the lowest fraction of isolates having high residuals and polynomial SVM model has the highest fraction of residuals. Sigmoid SVM has more uniform distribution of residuals than others. (d) Precision-recall curves. LR, LDA, and NBC outperform based on the residual distribution and precision-recall curve.

#### 4.4.5. bla_OXA-48_

According to Table 11, ML models containing Meropenem as one of their input features can be considered as the optimum antibiotic combination to predict *bla*_OXA-48_, given the simplicity principle and their performance. Considering the positive coefficient of Meropenem in the logistic function and sensitivity quantity, it can be concluded that if an isolate is resistant to the Meropenem antibiotic, it carries the *bla*_OXA-48_ gene with a probability of 56-94% with a 95% C.I. Also, following precision values in Meropenem-based models, 54-96% of the predictions indicating the presence of the *bla*_OXA-48_ gene in isolates are correct. In addition to classification metrics, the results of model-agnostic approach for feature selections are represented in Fig. 24. Moreover, residual distribution and precision-recall curve of all the algorithms based on the selected antibiotics through chi-squared test are shown in Fig. 25.

**Table 11.** Performance of ML Models predicting the presence of *bla*_*OXA-48*_ in the test dataset based on sensitivity, precision, and F1-Score.

| Feature Selection method | Features | LR |  |  | NBC |  |  | LDA |  |  | SVM-Sigmoid |  |  | SVM-Polynomial |  |  | CatBoost |  |  |
| --- | --- | --- | --- | --- | --- | --- | --- | --- | --- | --- | --- | --- | --- | --- | --- | --- | --- | --- | --- |
|  |  | S (95% C.I.) | P (95% C.I.) | F1 (95% C.I.) | S (95% C.I.) | P (95% C.I.) | F1 (95% C.I.) | S (95% C.I.) | P (95% C.I.) | F1 (95% C.I.) | S (95% C.I.) | P (95% C.I.) | F1 (95% C.I.) | S (95% C.I.) | P (95% C.I.) | F1 (95% C.I.) | S (95% C.I.) | P (95% C.I.) | F1 (95% C.I.) |
| Pearson's Chi-Squared test | AK+CTX+GN+IMI+FEP+CAZ+CIP+MEM+CRO+AZM | 0.65 (0.41-0.85) | 0.72 (0.47-0.90) | 0.68 (0.51-0.85) | 0.80 (0.56-0.94) | 0.59 (0.39-0.78) | 0.68 (0.53-0.83) | 0.80 (0.56-0.94) | 0.84 (0.60-0.97) | 0.82 (0.69-0.95) | 0.80 (0.56-0.94) | 0.84 (0.60-0.97) | 0.82 (0.69-0.95) | 0.65 (0.41-0.85) | 0.87 (0.60-0.98) | 0.74 (0.58-0.91) | 0.70 (0.46-0.88) | 0.74 (0.49-0.91) | 0.72 (0.56-0.88) |
| Wald test and Model agnostic in LR, NBC, CatBoost, SVM-Poly. & Sig. | IMI+MEM | 0.80 (0.56-0.94) | 0.84 (0.60-0.97) | 0.82 (0.69-0.95) | 0.80 (0.56-0.94) | 0.84 (0.60-0.97) | 0.82 (0.69-0.95) | 0.80 (0.56-0.94) | 0.84 (0.60-0.97) | 0.82 (0.69-0.95) | 0.80 (0.56-0.94) | 0.80 (0.56-0.94) | 0.80 (0.66-0.94) | 0.80 (0.56-0.94) | 0.80 (0.56-0.94) | 0.80 (0.66-0.94) | 0.80 (0.56-0.94) | 0.84 (0.60-0.97) | 0.82 (0.69-0.95) |
| Model agnostic in LDA | MEM | 0.80 (0.56-0.94) | 0.80 (0.56-0.94) | 0.80 (0.66-0.94) | 0.80 (0.56-0.94) | 0.80 (0.56-0.94) | 0.80 (0.66-0.94) | 0.80 (0.56-0.94) | 0.80 (0.56-0.94) | 0.80 (0.66-0.94) | 0.80 (0.56-0.94) | 0.80 (0.56-0.94) | 0.80 (0.66-0.94) | 0.80 (0.56-0.94) | 0.80 (0.56-0.94) | 0.80 (0.66-0.94) | 0.80 (0.56-0.94) | 0.80 (0.56-0.94) | 0.80 (0.66-0.94) |

MEM is applied to all five algorithms based on the simplicity principle to investigate their performance.

**Fig. 24.**
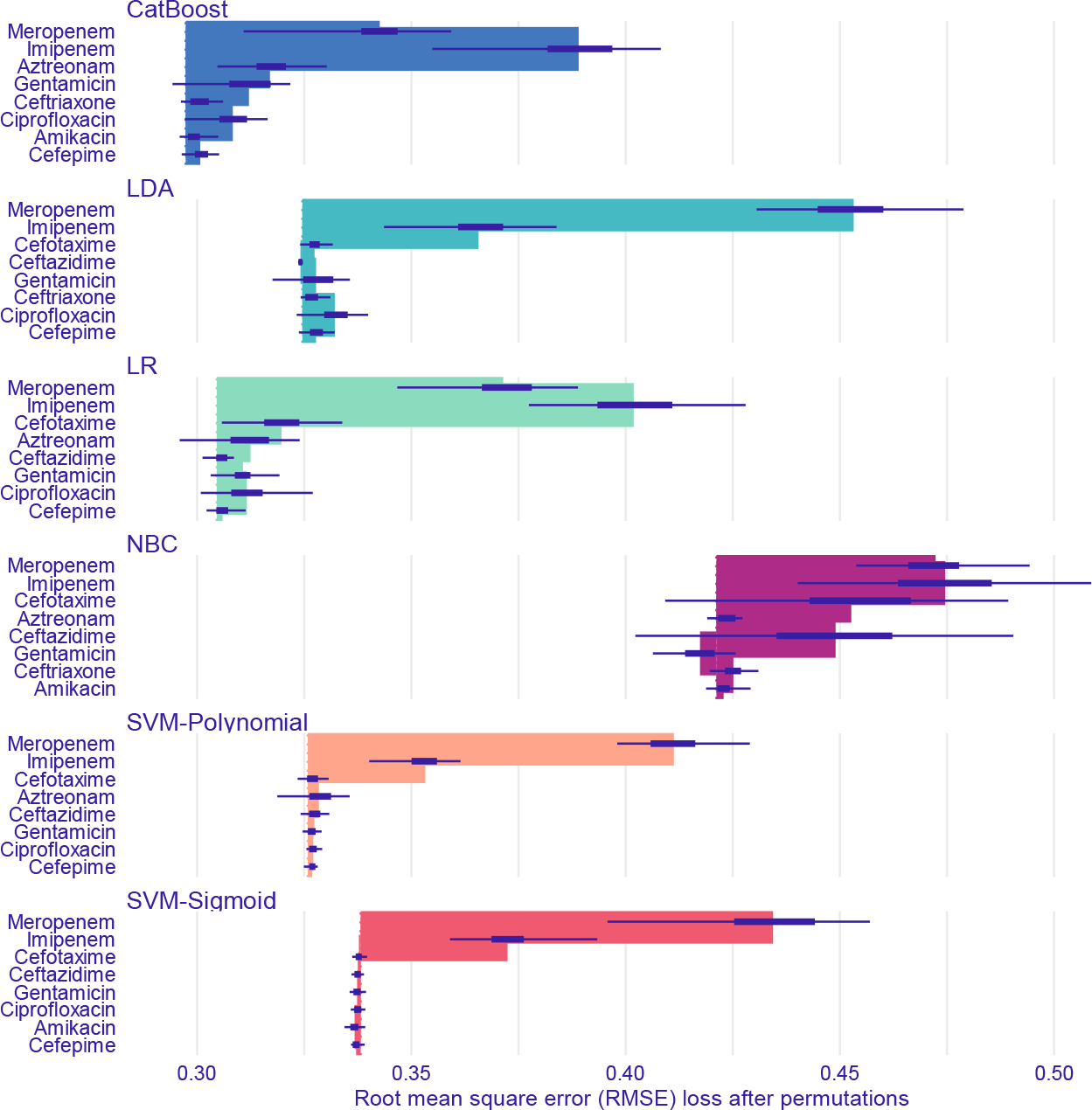
Feature importance of ML models predicting *bla*_OXA-48_ gene through model-agnostic approach over 50 permutations for top 8 antibiotics

**Fig. 25.**
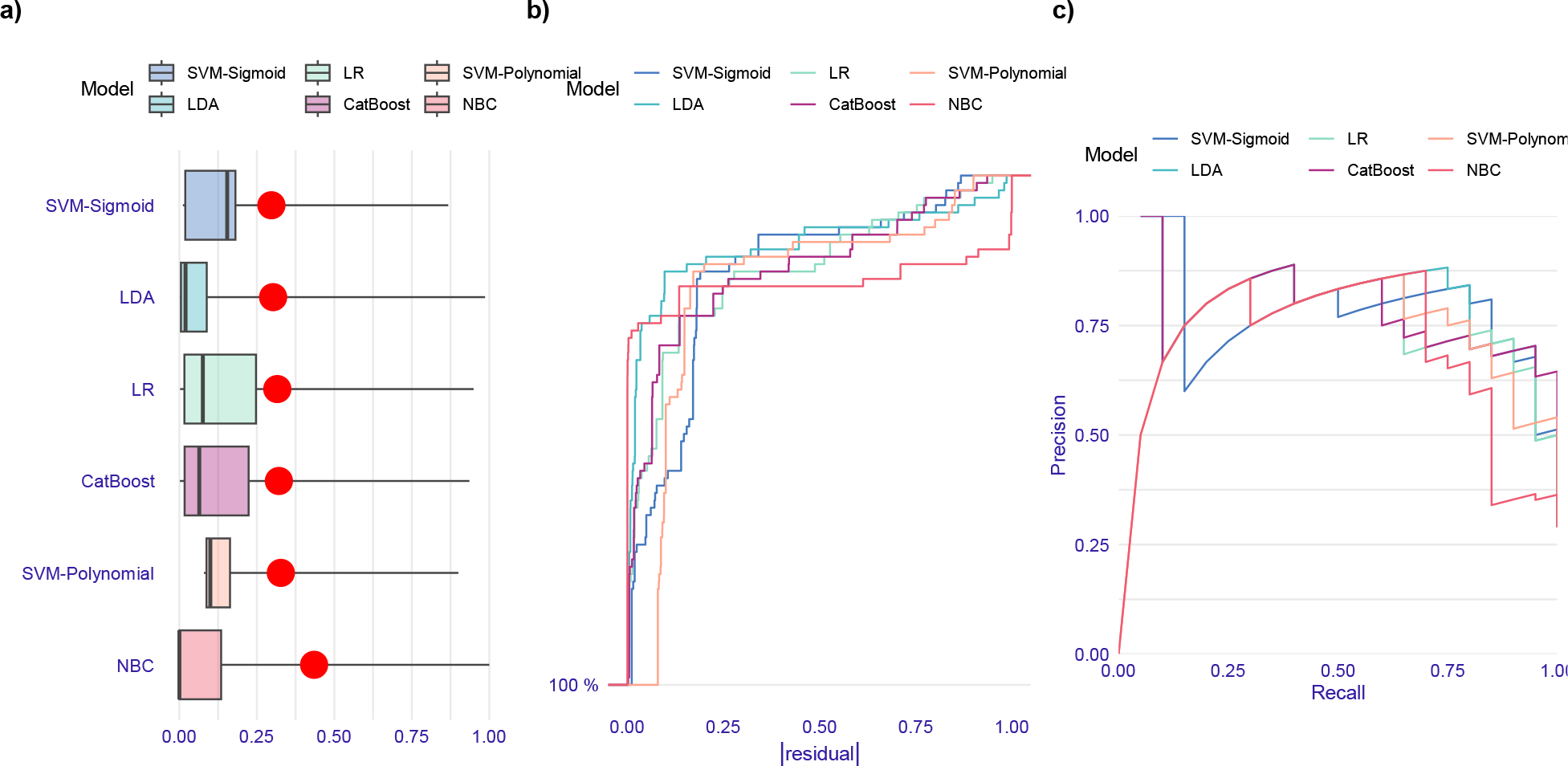
Evaluation of ML models performance in prediction of *bla*_OXA-48_ gene based on the test data set. (a) Boxplots of absolute values of residuals. Red dot stands for root mean square of residuals. (b) Histograms of residuals. (c) Reverse cumulative distribution of absolute values of residuals. Performance of all models, except NBC model, are close to each other. NBC model has the highest fraction of the isolates having high residuals. (d) Precision-recall curves. Sigmoid kernel of SVM and LDA models outperform others based on the residual distribution diagrams and precision-recall curve.

The OXA48-type carbapenemase was initially identified from carbapenem-resistant *K. pneumoniae* isolate in Turkey, it belongs to class D beta-lactamases and can hydrolyze penicillins, carbapenems and sparing expanded-spectrum cephalosporins (43, 44). In a another study conducted by Solgi *et al*., out of 71 isolates of carbapenem-resistant bacteria belonging to the *Enterobacteriaceae* family, 33 isolates (46.5%) carried the *bla*_OXA-48_ gene (Ref).

In our previous study reported in 2021, among 60 isolates of carbapenem-resistant hypervirulent *K. pneumoniae*, 45 isolates (75%) carried the *bla*_OXA-48_ gene (14). Therefore, it seems logical that among imipenem and meropenem-resistant isolates, the *bla*_OXA-48_ producing isolates are more prevalent, so there is a direct relationship between carbapenem resistance and the presence of the *bla*_OXA-48_ gene.

## 5. Discussion

*K. pneumoniae* has been mentioned as a bacterium that has experienced a significant increase in antibiotic resistance in the past decades, which has led to an increase in ineffective treatment, resulting in increased mortality and economic loss to healthcare systems (6). To detect antibiotic resistance routinely in microbiology laboratories, a culture-based antibiotic sensitivity test (AST) according to EUCAST or CLSI is used, which requires at least 48 hours, especially for samples like blood, enrichment is needed and requires more time, several rounds of culture, and also different antibiotic discs belonging to various classes to measure the bacterial antibiotics resistance. One of the most efficient ways to overcome infections caused by resistant bacteria is to rapidly identify their antibiotic resistance profile (45, 46). In order to optimize cost-efficiency and time savings, machine-learning techniques appear to be beneficial for predicting bacterial resistance. In this study, we evaluated and compared several machinelearning models to detect the presence of beta-lactamase and aerobactin genes and pathotypes. The ML models that we develop by the feature selection methods allow us to discover the influence of antibiotics on the performance indicators of each model. Moreover, considering signs of the feature coefficients (being positive or negative) in the logistic function, their impact (resistance or susceptibility) on predicting the presence of each gene and isolates’ pathotype identification could be determined. This will help us provide models using a limited number of antibiotics for discovering each gene and isolates’ pathotypes instead of analyzing all of them. To achieve this goal, we consider two conditions to select the most optimized model and antibiotic(s) combination:

1) For optimization purpose, classification metrics including accuracy, sensitivity, precision, and F1-Score should be maximized. However, a comprehensive maximization usually does not happen; some metrics will be more critical than others depending on the problem. For the current study, the algorithms’ ability to detect genes’ presence and isolates’ pathotypes is more critical than determining their absence, therefore, optimization of the criteria focusing on the isolates carrying beta-lactamase and aerobactin genes and pathotypes is a main priority. Maximized sensitivity increases the probability of the classifier’s success in detecting isolates carrying genes. In addition, the algorithm’s predictions representing the presence of genes/pathotypes should be as accurate as possible. That means the high performance of the algorithm in the prediction process concerning precision value. Due to considering the precision and the sensitivity metrics, the F1-score is employed to provide a trade-off between them. As a result, ML models with a specific antibiotic(s) combination perform better than the others, if they have higher sensitivity and precision, consequently, F1-score. Otherwise, the ML model with the antibiotic(s) combination which led to higher sensitivity but close precision and F1-score compared to other models and combinations could be selected as the best model and antibiotic(s) combination for correct detection of the pathotype/gene.

In addition to the conventional metrics (e.g., sensitivity, precision, accuracy, and F1-score), we performed more analyses through boxplots of residuals, precision-recall curves, and reverse cumulative distribution (RCD) curve (47) of residuals. These analyses are useful especially when the models seem to perform very similarly based on the metrics extracted through confusion matrix. Therefore, these analyses provide a clear intuitive illustration so that help us make a better comparison between the performances of models. The RCD curve of residuals represents the percentage of isolates having at least a specific absolute residual. Each point on the curve shows the percentage of isolates having equal to or greater than a specified absolute value of residual. RCD plots alongside boxplots of residuals give more precise perspective about distribution of residuals and their relationships can be traceable by comparison with each other.

2) There is a trade-off between the interpretability and accuracy of the model (48). According to the simplicity principle, if the simpler model’s performance and prediction ability are similar to the more complex model (a model with more features), the former model’s interpretability and practical conclusions outweigh the latter, so the simpler model is preferable. The prediction performance will increase with the complexity of the model, but that will boost the difficulty of interpreting the result. Also, there is a higher risk of over fitting in complex models. Therefore, considering these issues leads us to select simpler models if there is no noticeable decrease in metrics, especially in the three mentioned metrics in the first part (28, 49).

## 6. Concluding remarks

The proposed approach in this study specifies the most effective antibiotics that helps identify pathotypes, aerobactin and beta-lactamase genes in experimental applications by methodologically selecting crucial antibiotics that results in less number of required experiments on more limited antibiotics. Implementing a machine learning approach using various feature selection methods and predictive algorithms for detecting different pathotypes and genes results in specifying more limited significant antibiotics for antimicrobial susceptibility testing. Moreover, in the case of detection of the hv*Kp* strain directly by ML, we could identify a limited number of antibiotics for detecting the hv*Kp* pathotype without investigating *iutA* and *iucA* genes. For instance, Cefepime, as a common crucial antibiotic in detecting both *iutA* and *iucA* genes, is selected by ML models as an ineffective antibiotic for detecting hv*Kp*.

Finally, we claim that our proposed approach can be used as a fast affordable and reliable way that can significantly reduce the work-load in detection of classical and hypervirulent pathotypes and beta-lactamase and aerobactin genes.

## Code and data availability

All codes (written in R) and data are available through Github: https://github.com/kouroshAKiani/ML-GeneticPathotyping

## Appendix

### Logistic Regression

The Logistic regression (LR) algorithm is a statistical method for modeling the probability of occurrence of an event using the Logistic function (19).

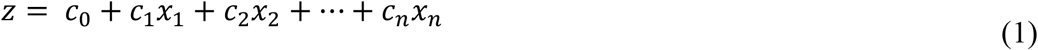

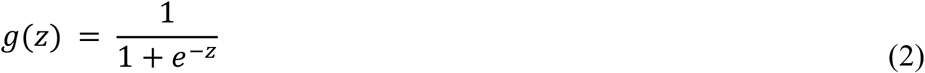

in which *g*(*z*) is the Logistic function, and *Z* is a linear combination of dependent variables. The Logistic function’s output is a value between 0 and 1, which is the probability of occurrence of an event.

*g*(*z*) Means *P*(*x* = *X*), i.e., the probability of predicted events and the likelihood of predicted non-events is *P*(*x* = *X*) = 1 − *h*(*c*).

### Naïve Bayes Classifier

Naïve Bayes Classifier (NBC) is a simple probabilistic classifier based on applying ‘Bayes’ theorem with independence assumptions between predictors, that is 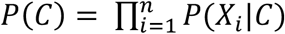, where *X* = (*X*_1_, …, *X*_*n*_) is a feature vector, and C is a class (24). There are three hyperparameters (50) in the NBC algorithm that are tuned through 10 fold cross-validation:

1) Type of density estimate; specifies kernel density estimate or Gaussian density estimate.

2) Kernel density bandwidth; more extensive bandwidth results in a more flexible density estimate.

3) Laplace smoother; ensures that the probability of occurring for each class in each feature is nonzero.

### Linear Discriminant Analysis

Linear discriminant analysis (LDA) is a well-established statistical-based pattern classification method for finding linear combinations of variables that best separate observations into groups or classifications. The LDA method identifies a projection vector for two-class problems to maximize the between-class and within-class variance ratios (25).

### Support Vector Machines

Support vector machines (SVMs) are learning algorithms introduced initially by Vpanik and co-workers (27). We define a hyperplane as the classifier in the SVM algorithm to classify the data with different classes. Various kernels, such as the sigmoid kernel, Radial Basis Function (RBF) kernel, polynomial kernel, and linear kernel, could be used in SVMs. The “C” parameter is the most crucial variable in the SVM algorithm, and its precise tuning greatly influences the model’s performance. The “C” parameter determines the margin of the developed hyperplane from the examples of each class label. The goal of “C” parameter tuning is that the hyper-plane separates the classes as accurately as possible. Two margins for hyperplane could be considered:

A) Large (C) – maximal margin

B) Soft (C) – soft margin

The maximal margin at the training stage significantly decreases the learner error (minimizing bias), but minimized bias results in overfitting and increases the algorithm’s error in the test stage (maximizing variance). Therefore, we control the bias-variance trade-off of the statistical learning via the C parameter, which is inversely proportional to regularization. Optimized hyper-parameters of each kernel and the “C” parameter will be tuned through 10-fold crossvalidation.

#### 3.2.4.1 Polynomial kernel function

Eq. 3 is the governing function of the Polynomial kernel function (51);

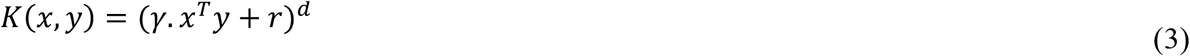

γ, r, and d are hyper-parameters tuned for each model using 10-fold cross-validation.

#### 3.2.4.2 Sigmoid kernel function

Eq. 4 is the governing function of the Sigmoid kernel (51);

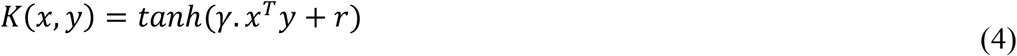

in which γ and *r* are the hyper-parameters of this kernel that are tuned through 10-fold crossvalidation.

### CatBoost

CatBoost is an improved gradient-boosting algorithm proposed in 2018 (26). CatBoost tackles classification and regression problems and has been publicized in an open-source multiplatform gradient-boosting library (26, 52). Decision trees are employed as a weak base learner in the CatBoost algorithm and gradient boosting to fit the decision tree sequentially. CatBoost is a large-scale and comprehensive library composed of various elements such as GPU learning and standard boosting and consists of 10-fold hyperparameters optimization to modify to several practicable investigation circumstances. In this study, four essential hyperparameters (26) were tuned through 10-fold cross-validation for the CatBoost algorithm, including tree depth, L2 regularization coefficient, learning rate, and iterations.

Based on the resistance or susceptibility of isolates to the antibiotics, several predictive models by machine learning algorithms will be trained. Hyperparameters (if applicable) will be tuned by employing 10-fold cross-validation. Then, the test data will assess each model’s performance and generalization ability. There are various criteria for classification problems to evaluate the model’s performance. The quantity of these criteria can be obtained by using a confusion matrix.

### Confusion matrix

Different parameters like sensitivity, specificity, accuracy, precision, and F1-score (53-55) are calculated to measure the performance and generalization ability of the ML classifier, as follows:

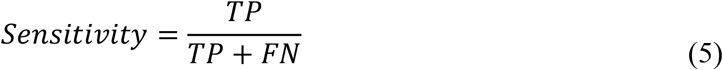

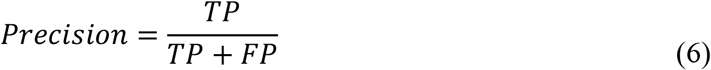

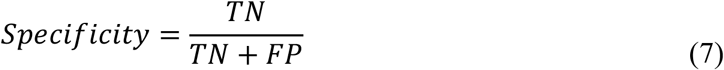

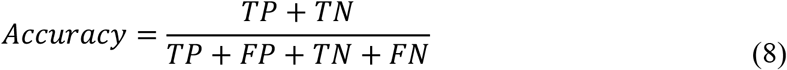

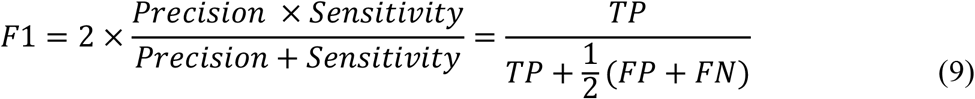

Where TP is defined as true positive, representing the total number of correctly predicted isolates carrying a beta-lactamase or belonging to a strain, TN is defined as True Negative, which means the total number of correctly predicted isolates not carrying a gene or not belonging to a strain. FP means False Positive, which expresses the total number of isolates not carrying a gene or not belonging to a pathotype classified as isolates carrying a gene or belonging to a strain. In contrast, FN means False negative, which defines the total number of isolates carrying a gene or belonging to a strain classified as isolates not carrying a gene or not belonging to a strain (53). Applying the trained model to isolates available in the dataset will make a prediction that should be verified according to the actual data. Therefore, the confusion matrix will be utilized to assess all the classification models’ performances in the test stage.

